# Neuronal interactions in forebrain organoids lead to protective antiviral responses

**DOI:** 10.1101/2024.05.07.592842

**Authors:** Seble G. Negatu, Christine Vazquez, Carl Bannerman, Kevin R. Amses, Guo-li Ming, Kellie A. Jurado

## Abstract

Neurotropic viruses are the most common cause of infectious encephalitis and largely target neurons for infection. Our understanding of intrinsic neuronal innate immune response capacity to mediate protective antiviral responses remains incomplete. Here, we evaluated the role of intercellular crosstalk in mediating intrinsic neuronal immunity and its contribution to limiting viral infection. We found that in the absence of viral antagonism, neurons transcriptionally induce robust interferon signaling and can effectively signal to neighboring uninfected bystander neurons. Yet, in two-dimensional cultures, this dynamic response did not restrict viral spread. Interestingly, this differed in the context of viral infection in three-dimensional forebrain organoids with complex neuronal interactions, where we observed protective capacity. We show antiviral crosstalk between infected neurons and bystander neural progenitors is mediated by type I interferon signaling. Using spatial transcriptomics, we then uncover distinct regions of bystander progenitor interactions that reveal critical underpinnings of protective antiviral responses, including expression of distinct antiviral genes. These findings underscore the importance of intercellular communication in protective antiviral immunity in the brain and implicate key contributions to protective antiviral signaling.

## INTRODUCTION

The brain houses a mosaic of molecularly distinct neuronal subtypes that form complex intercellular networks. Neurotropic viruses target the brain for infection, often with a preference for replication within neurons. Neuronal subtypes have differential innate immune signatures that are influenced by regionality or maturation state.^1, 2^ Some studies suggest that neurons are capable of sensing pathogens and inducing responses, while others propose that neurons have high viral burdens due to the absence of an effective antiviral response.^3–7^ These seemingly conflicting results indicate that our understanding of intrinsic neuronal innate immune responses is incomplete.

The type I interferon (IFN) response is the primary driver of an antiviral state in many cells. To evade activation of innate immune responses, many viruses have evolved to antagonize type I IFN signaling. For example, La Crosse virus (LACV) is a neuron-targeting RNA virus that is the primary cause of pediatric encephalitis in the United States, which has a non-structural protein in its S segment (NSs) that blunts antiviral IFN responses by broadly inhibiting host cell transcription.^8, 9^ Viruses antagonize immune responses in infected cells, which makes immune signaling in uninfected bystander cells essential for limiting viral spread. The interplay of immune responses between uninfected bystander and infected neurons remains unexplored.

This gap led us to investigate the role of intercellular crosstalk in mediating intrinsic neuronal immunity and its contribution to protective antiviral responses. We demonstrate that neurons have a strong ability to sense viral infection and elicit robust innate immune signatures when viral antagonism is limited. Using a forebrain organoid model with complex neuronal interactions, we reveal antiviral crosstalk between infected neurons and bystander neural progenitors that is mediated by type I IFN signaling. Using spatial transcriptomics, we then uncover distinct regions of bystander progenitor interactions that reveal critical underpinnings of protective antiviral responses in neurons. These findings underscore the importance of intercellular communication in protective antiviral immunity in the brain and implicate key contributions to protective antiviral signaling.

## RESULTS

### Viral antagonism masks neuronal intrinsic capacity to induce robust interferon signaling

Neurons have long been considered poor IFN producers; however, neuronal potential to mount effective IFN responses to viral infection has been primarily modeled using viruses that antagonize innate immune activation.^10^ Thereby, we first evaluated if neurons increased capacity to intrinsically induce antiviral pathways in the absence of immune antagonism. We cultured embryonic murine cortical neurons for 9-days prior to infection with WT-LACV or ΔNSs-LACV, a strain that lacks the well-defined viral antagonist NSs (ΔNSs-LACV mutant validation in **Supplemental Figures 1A** and **1B**).^9, 11^ Additionally, we exposed neuronal cultures to heat- or UV-inactivated ΔNSs-LACV to evaluate neuronal responses to viral pathogen associated molecular patterns (PAMPs) in the absence of viral replication. To define neuronal innate immune potential in an unbiased manner, we conducted bulk RNA sequencing at 16-hours post-infection (HPI) (**Figure 1A**). WT- and ΔNS-LACV-infected neurons formed distinct clusters on a PCA plot indicating they had unique responses to infection (**Figure 1B**). In contrast, neuronal cultures treated with heat- or UV-inactivated ΔNSs-LACV largely clustered with mock, signifying viral replication was essential to drive robust changes in neuronal gene expression.

**Figure 1:**
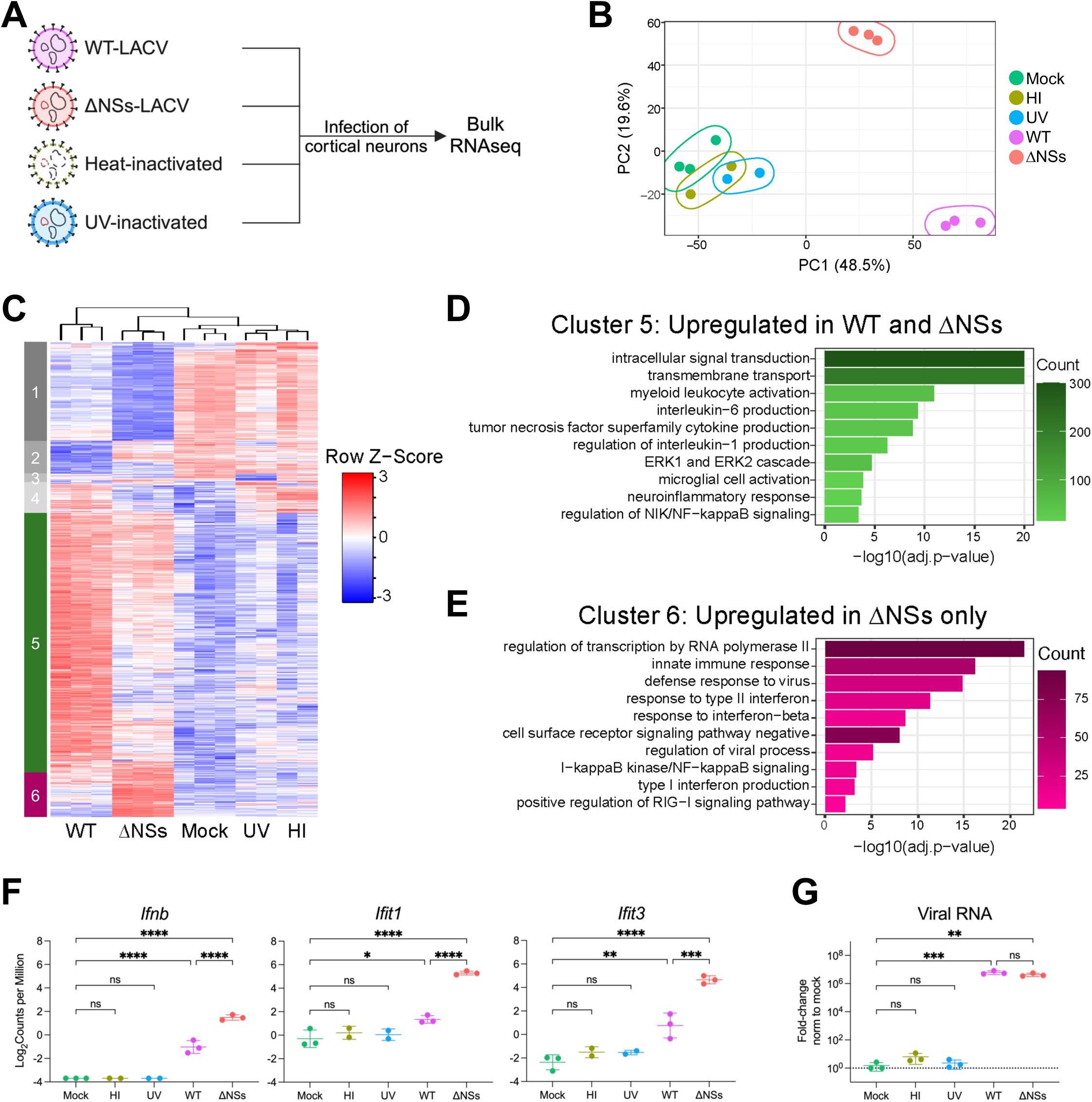
Viral antagonism masks neuronal intrinsic capacity to induce robust interferon signaling. (A) Schematic of experimental design indicating primary murine cortical neurons that were infected at 0.5 MOI for 16 hours with wild-type La Crosse Virus (WT-LACV), recombinant ΔNSs La Crosse Virus (ΔNSs-LACV), heat-inactivated (HI) ΔNSs-LACV, or ultraviolet (UV)-inactivated ΔNSs-LACV followed by bulk RNA sequencing of total RNA. (B) Principal component analysis (PCA) showing PC1 and PC2 for bulk RNA-seq data from infected neurons. (C) Heat map of expression bulk RNA sequencing. Columns represent samples clustered using Spearman correlation. Pearson correlation was used to cluster differentially expressed genes (cutoffs: p-value=0.01 and log fold-change= ±1) compared to mock, which were then represented as heatmap with the data scaled by z-score for each row. (D-E) Functional enrichment analysis of genes from cluster 5 (D) and cluster 6 (E) using Gene Ontology biological process. (F) Log2 adjusted counts per million of select innate antiviral genes. Mean±SD. Neurons pooled from 2 litters. One-way ANOVA followed by Tukey’s test. (G) Fold change of LACV RNA as determined by quantitative RT-PCR relative to the housekeeping gene *hprt*, normalized to mock. Neurons pooled from 2 litters. Mean±SD. One-way ANOVA followed by Tukey’s test, ns p>0.05, *p<0.05, **p<0.01, ***p<0.001, and ****p<0.0001.

Unbiased clustering of 4057 differentially expressed genes revealed 6 unique clusters (**Figure 1C**). Cluster 5 represented genes upregulated in both WT- or ΔNSs-LACV infected neurons but was more highly induced with WT-LACV infection. Pathway analysis of these genes uncovered activation of inflammatory pathways such as NF-κB signaling and production of cytokines including tumor necrosis factor (**Figure 1D**). In contrast, cluster 6 denoted genes that were uniquely upregulated in the ΔNSs-LACV infected samples. Regulation of transcription by RNA polymerase II was upregulated in ΔNSs-LACV-infected samples, corroborating that the NSs protein antagonizes host gene expression by limiting RNA polymerase II function (**Figure 1E**). Interestingly, functional enrichment analysis showed specific induction of viral recognition machinery such as RIG-I signaling and the type I IFN antiviral pathway in the absence of viral antagonism. To evaluate drivers of this specificity in ΔNSs-LACV-infected neurons, we queried representative innate antiviral genes in cluster 6. We observed increased gene expression of upstream signaling mediators such as *Ifnb* and downstream IFN-stimulated genes (ISGs) including *Ifit1* and *Ifit3* in ΔNSs-LACV-infected neurons compared to WT-LACV (**Figure 1F**). These differences are not attributable to viral PAMP level since WT- and ΔNSs-LACV had similar viral RNA levels (**Figure 1G**). These data suggest that in the absence of viral antagonism, primary murine cortical neurons have a strong capacity to sense pathogen and induce expression of IFN-specific genes.

### Uninfected bystander neurons induce innate immune responses

Given that ΔNSs-LACV infection of murine neurons distinctly elicited robust IFN signaling, we used it to investigate whether the observed antiviral responses were unique to infected neurons or nearby uninfected bystander neurons. We utilized validated LACV-targeting fluorescence in-situ hybridization (FISH) probes (**Supplemental Figure 1C**) to demarcate ΔNSs-LACV-infected neurons prior to fluorescence-activated cell sorting (FACS) for bulk RNA sequencing (**Figure 2A**).^12, 13^ We used mock-treated neurons as a threshold to define uninfected bystander cells within infected cultures, and positive LACV FISH signal to identify highly infected neurons. The remaining cells were deemed an intermediately infected population. To identify a timepoint where both uninfected bystander and highly infected populations are distinguishable, neurons from infected cultures were collected at 1-, 16- and 24-HPI for FISH staining followed by flow cytometry (**Figure 2B**). Quantification of the bystander, intermediate and highly infected neurons revealed a shift from majority bystanders at 1-HPI to highly infected neurons by 24-HPI (**Figure 2C**). At 16-HPI we observed a bimodal distribution enabling clear separation of bystander and highly infected neurons. To validate sorting at this timepoint, we FACS-sorted ΔNSs-LACV-infected neurons at 16-HPI and visualized LACV FISH signal using fluorescence microscopy (**Figure 2D**). As suggested by the bimodal distribution of bystander and highly infected neurons seen by flow cytometry, uninfected bystanders lacked FISH signal, similar to mock, while highly infected neurons were all positive. However, neurons in the intermediate population had a mixed positive and negative LACV FISH signal, leading us to omit this group from further analysis.

**Figure 2:**
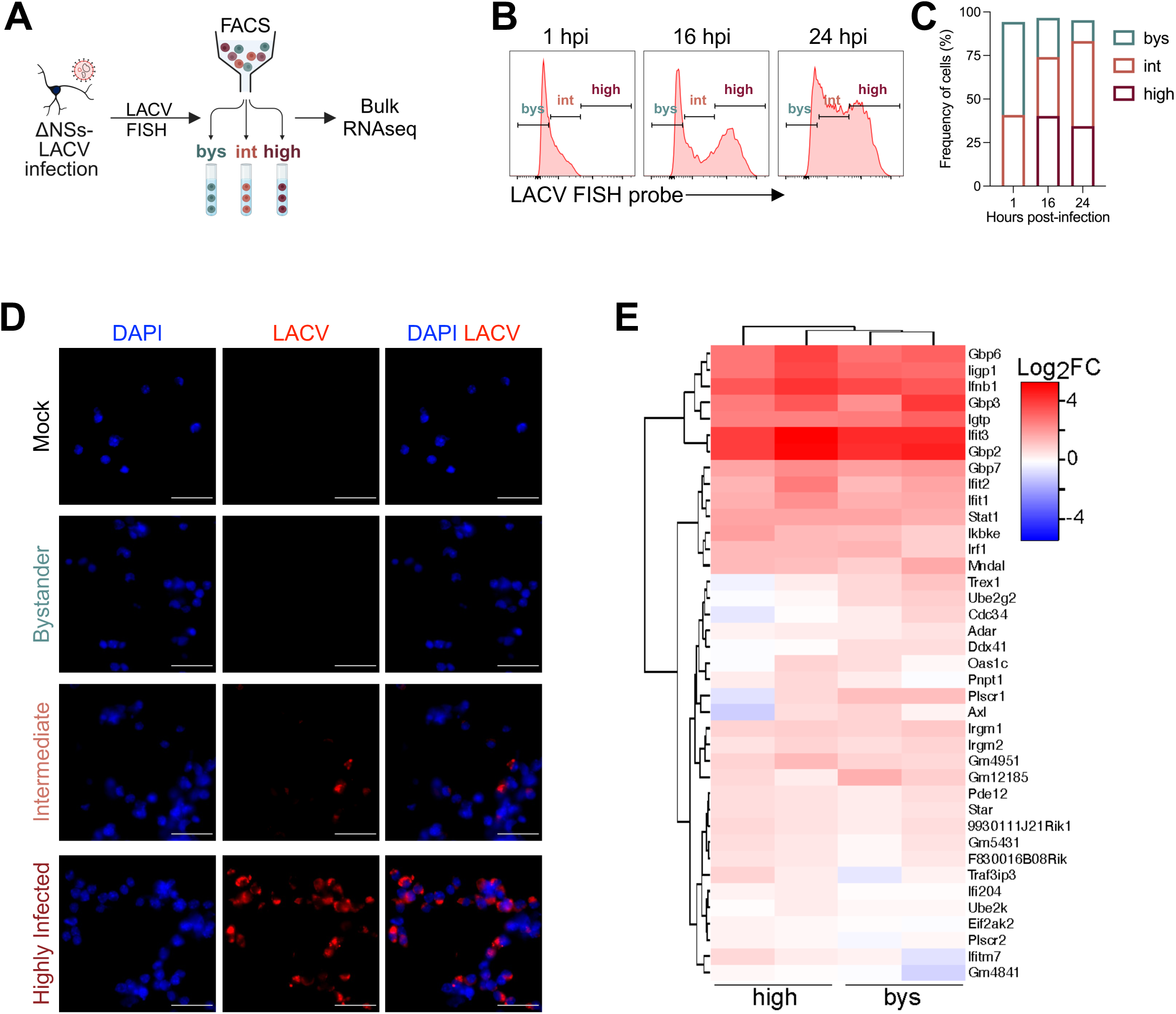
Uninfected bystander neurons induce innate immune responses. (A) Schematic of adapted probe-seq pipeline where murine cortical neurons are infected with ΔNSs-LACV, then stained with LACV FISH probes prior to FACS sorting of three populations, bystander (bys), intermediate (int) and highly infected (high) neurons, for bulk RNA sequencing. (B-C) Gating (B) and quantification (C) of neurons at 1-, 16- and 24-hours post-infection (HPI). Neurons pooled from 1 litter. (D) Images of FACS-sorted ΔNSs-LACV -infected neurons at 16-HPI. LACV RNA labeled by FISH probes (red); cell nuclei (blue). Scale bars, 25μm. (E) Heatmaps of log_2_ fold-change of select antiviral genes using unsupervised clustering. Columns represent infected (high) and bystander (bys) neurons relative to mock-treated cells. Selected genes were derived from MSigDB GoBP: Response to interferon alpha and beta gene sets. Neurons pooled from 1 litter per independent experiment; N=2.

For sequencing analysis, RNA was extracted from sorted uninfected mock, uninfected bystander and infected populations. Given the specific innate antiviral gene signature we observed in ΔNSs-LACV-infected neurons (**Figure 1E**), we sought to compare the expression of type I IFN response genes between uninfected bystander and infected populations. Expression of genes selected from curated datasets of the IFN alpha/beta response pathway were evaluated (**Figure 2E**). Surprisingly, unsupervised clustering revealed similar ISG expression patterns by both bystander and highly infected neurons, with the *Ifit* family of genes having elevated expression.

Taken together, these data show that during ΔNSs-LACV infection of neurons, uninfected bystanders that neighbor infected cells contribute to the robust type I IFN transcriptional signature observed. However, despite these differences, both viruses exhibited nearly identical growth curves (**Supplemental Figures 1D** and **1E)**, indicating that the transcriptional induction of ISGs during ΔNSs-LACV infection does not limit viral replication or virion production in this model system.

### Interneuronal networks within human forebrain organoids reveal protective ISG production by bystander neural progenitors

To better model the intercellular neuronal network and complexity of neuronal antiviral responses, we employed human induced pluripotent stem cell (iPSC)-derived forebrain organoids which mimic features of the developing human brain. This three-dimensional architecture and heterogeneity of neuronal differentiation states leverages our ability to assess complex intercellular interactions that contribute to innate immune responses.

To first model the radial organization of progenitors, we generated 35-days *in-vitro* (DIV) forebrain organoids which contain neural progenitor regions demarcated by SOX2 expression, commonly referred to as neural rosettes (**Supplemental Figure 2A and 2B**). We then infected with WT- or ΔNSs-LACV for 24 hours and collected samples at 4-days post-infection (DPI) (**Figure 3A**). At 4-DPI, we saw significantly less viral RNA during ΔNSs-LACV infection as compared with WT-LACV (**Figure 3B**). Furthermore, *IFIT1* gene expression was lower in mock-treated organoids, and higher in ΔNSs-LACV infected forebrain organoids compared to those infected with WT-LACV (**Figure 3C**). Together, this suggests the potential of forebrain organoids to elicit an effective antiviral response in the absence of immune antagonism.

**Figure 3:**
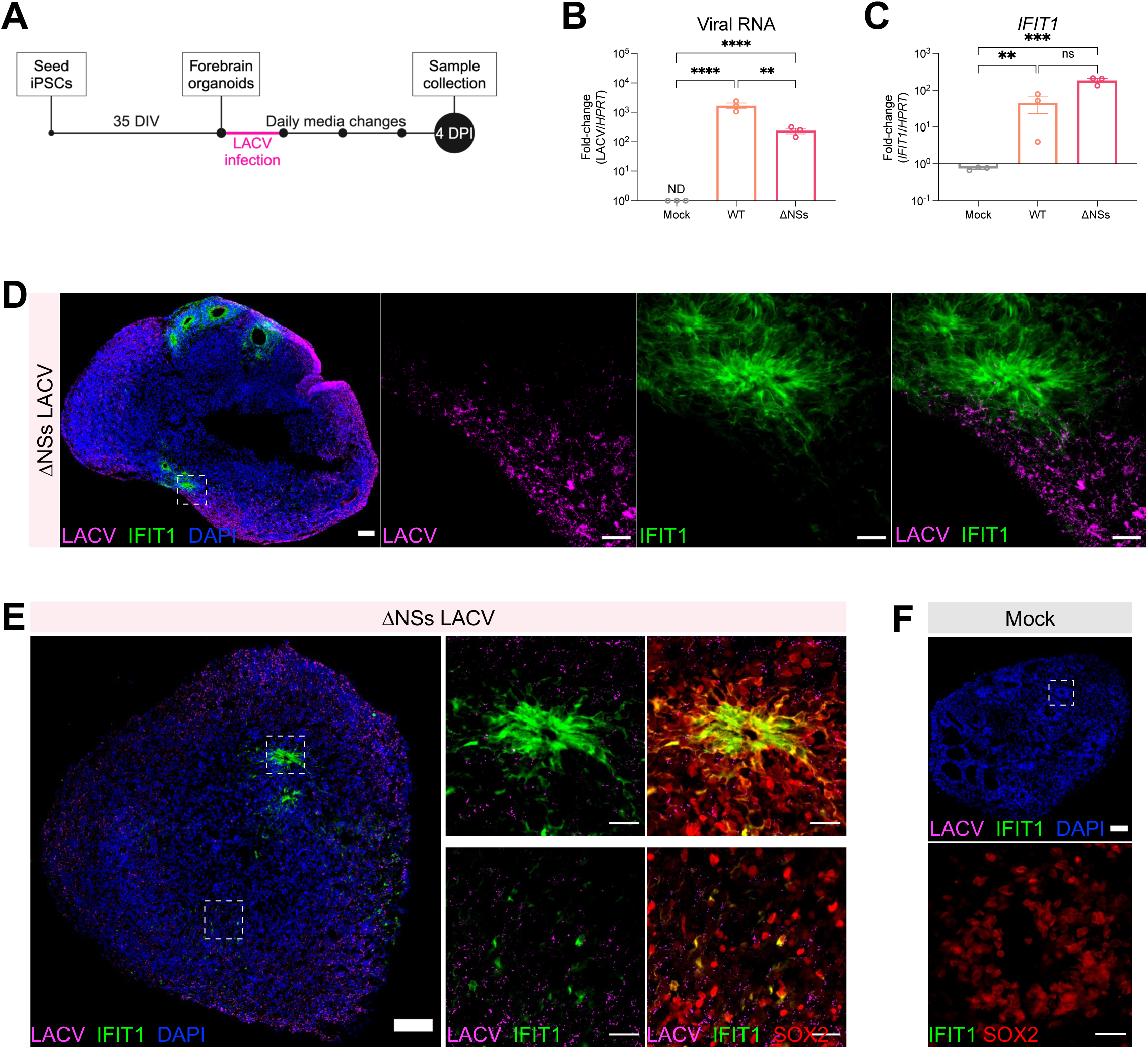
Interneuronal networks within human forebrain organoids reveal protective ISG production by bystander neural progenitors. (A) Schematic of experimental design depicting 35-days *in-vitro* (DIV) human induced pluripotent stem cell-derived forebrain organoids infected with 2.5×10^6^ PFU WT-LACV, ΔNSs-LACV, or mock treated for 24 hours followed by sample collection at 4-days post-infection (DPI). (B-C) Fold change of LACV RNA (B) and *IFIT1* (C) was determined by quantitative RT-PCR relative to the housekeeping gene *HPRT*. N=3. Mean±SEM. One-way ANOVA followed by Tukey’s test, ns p>0.05, **p<0.01, ***p<0.001, ****p<0.0001. (D) Immunofluorescent images of ΔNSs-LACV infected organoids. White box indicates inset. LACV glycoprotein (magenta); IFIT1 (green); cell nuclei (blue). Scale bars, zoom-out 100μm, inset 25μm. (E-F) Immunofluorescent images of ΔNSs-LACV-infected (E) or mock-treated (F) organoids. White dashed boxes indicate insets. LACV glycoprotein (magenta); IFIT1 (green); Sox2 progenitor (red); cell nuclei (blue). Scale bars, zoom-out 100μm, inset 25μm. Representative of 3 independent samples.

To visualize virus localization and IFIT1 protein expression in the absence of viral antagonism, we collected ΔNSs-LACV-infected forebrain organoids for immunofluorescence at 4-DPI (**Figure 3D**). We detected substantial viral antigen throughout the periphery of the organoids. Furthermore, IFIT1 expression radiated from neural rosettes, regions which lacked LACV signal, suggesting that ISGs were derived from bystander cells. Given the radial IFIT1 expression pattern, we sought to further characterize IFIT1+ cells by staining ΔNSs-LACV-infected (**Figure 3E**) and mock-treated organoids (**Figure 3F**) for SOX2 expression, a neural progenitor marker. We observed colocalization of IFIT1 expression and SOX2 within and outside of neural rosettes (**Figure 3E**). Mock-treated organoids lacked basal IFIT1 protein expression, despite the presence of SOX2+ neural progenitors (**Figure 3F**), suggesting a viral-driven IFIT1 induction.

Overall, we found that human forebrain organoids can serve as a model for neuronal-intrinsic antiviral responses. Specifically, we observed that uninfected progenitors induced innate signaling that limited viral replication, suggestive that bystander neural progenitors play a role in antiviral protection.

### Interferon signaling within bystander neural progenitors limits viral spread and replication

We next sought to investigate the mechanism by which bystander neural progenitors contribute to the antiviral response in human forebrain organoids. First, to determine whether the innate immune responses we observed occurred through canonical IFN pathways, we used a janus-kinase (JAK) inhibitor, ruxolitinib, which blocks JAK1 and JAK2 downstream components of IFN signaling. Specifically, we infected 35-DIV forebrain organoids with ΔNSs-LACV for 24 hours and sustained ruxolitinib treatment until 4-DPI for collection and immunofluorescence staining (**Figure 4A**). JAK inhibition promoted viral antigen spread toward the center of the organoid, whereas intact JAK signaling during ΔNSs-LACV infection alone limited virus to the periphery. Quantification of viral staining indicated significantly more ΔNSs-LACV signal dissemination with ruxolitinib treatment (**Figure 4B**). This was further supported by qPCR for viral RNA (**Figure 4C**). Moreover, inhibition of JAK signaling completely eliminated IFIT1 expression in bystanders as determined by immunofluorescence and qPCR (**Figure 4D and 4E**), indicating a lack of bystander innate activation with ruxolitinib treatment. This reveals that intrinsic JAK-specific/dependent neuronal responses are capable of limiting viral spread in forebrain organoids.

**Figure 4:**
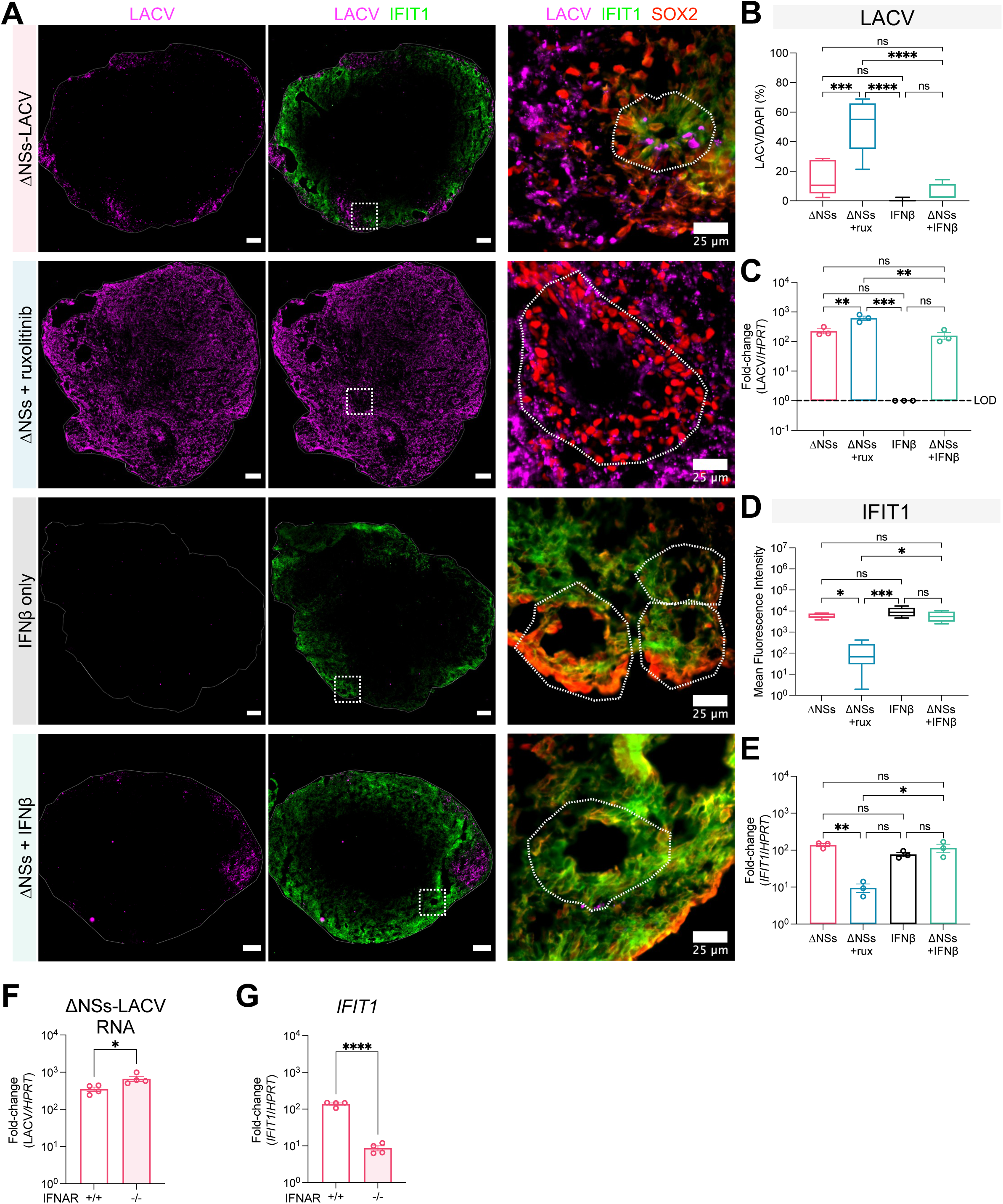
Interferon signaling within bystander neural progenitors limits viral spread and replication. (A) Immunofluorescent images of 35-DIV forebrain organoids infected with 2.5×10^6^ PFU ΔNSs-LACV alone; ΔNSs-LACV with 20uM ruxolitinib (janus-kinase inhibitor); treated with 100 units/mL IFNβ alone; or ΔNSs-LACV with 100 units/mL recombinant IFNβ. Infection for 24 hours followed by daily media changes and sustained ruxolitinib or recombinant IFNβ treatment until sample collection at 4-DPI. White dashed boxes indicate insets. LACV glycoprotein (magenta); IFIT1 (green); SOX2 progenitor (red); cell nuclei (blue). Scale bars, zoom-out 100μm, inset 25μm. (B) Upper, quantification of LACV+ area relative to DAPI+ area in immunofluorescent images (A). N=3, with 2-3 sections per experiment. Bar represents mean and whiskers represent ± SEM. Lower, fold change of LACV RNA as determined by quantitative RT-PCR relative to the housekeeping gene *HPRT*. N=3. Mean±SEM. One-way ANOVA followed by Tukey’s test. (C) Upper, quantification of IFIT1 mean fluorescent intensity in immunofluorescent images (A). N=3, with 2-3 sections per experiment. Bar represents mean and whiskers represent ± SEM. Lower, fold change of *IFIT1* as determined by quantitative RT-PCR relative to the housekeeping gene *HPRT*. N=3. Mean±SEM. One-way ANOVA followed by Tukey’s test. (D-F) WT and IFNAR1 knockout forebrain organoids infected with 2.5×10^6^ PFU ΔNSs-LACV for 24 hours followed by daily media changes. (D) Immunofluorescent images of infected WT and IFNAR knockout organoids. LACV glycoprotein (magenta); IFIT1 (green). Scale bars, 100μm. Fold change of LACV RNA (E) and *IFIT1* (F) as determined by quantitative RT-PCR relative to the housekeeping gene *HPRT*. N=5. Mean±SEM. Unpaired Student’s t test, ns p>0.05, *p<0.05, **p<0.01, ***p<0.001, ****p<0.0001.

Next, to investigate if IFIT1 expression in bystanders is mediated by type I IFN, we treated forebrain organoids with recombinant IFNβ to activate type I IFN independent of viral infection. *IFIT1* expression was upregulated along the periphery; however, both *SOX2* positive and negative cells expressed *IFIT1* (**Figure 4A**), indicating that innate immune activation was no longer limited to progenitors. Treatment with recombinant IFNβ shows that both progenitors and non-progenitors have capacity for innate signaling. To see if this broad innate activation better restricts viral spread, we treated forebrain organoids with recombinant IFNβ throughout infection. We observed virus localization and viral RNA levels similar to infection alone (**Figure 4A and 4B**). Moreover, sustained IFNβ treatment throughout infection led to a similar induction of IFIT1 compared to infection alone (**Figure 4A and 4C**). These data indicate that intercellular communication during viral infection sufficiently activates protective type I IFN pathways.

To directly evaluate the role of type I IFN in this intrinsic innate response, we next used CRISPR/Cas9 to knock-out the type I IFN receptor, IFNAR1, in iPSCs to generate IFNAR1 knockout forebrain organoids (**Supplemental Figure 3**). Here, we observed increased viral RNA in the IFNAR1 knockouts (**Figure 4E**) and diminished *IFIT1* expression (**Figure 4F**), indicating a direct role for type I IFN signaling. Cumulatively, these data indicate that innate immune activation by bystander neural progenitors contribute to the antiviral response in human forebrain organoids. Specifically, neuronal interactions between infected and bystander cells limit viral spread through type I IFN-dependent intercellular communication.

### Spatial transcriptomics uncovers distinct regions of progenitor bystander activation

We leveraged spatial transcriptomics to investigate the interplay of bystander activation and protection during infection in forebrain organoids. Using the Visium platform, we obtained transcriptomes of 1409 spots in 8 ΔNSs-LACV-infected organoids and 1478 spots in 9 mock-treated organoids (**Supplemental Figure 4A and 4B**). Each individual spot is 55μm and thereby contains several cells.^14^ With the goal of understanding neuronal interactions with respect to regions of viral infection, we aligned spatial data to LACV-stained organoids. We then conducted a clustering analysis that revealed seven distinct clusters (**Figure 5A and 5B**). Each cluster represents a different organoid region, which are all represented in both ΔNSs-LACV-infected and mock-treated conditions (**Supplemental Figure 4C**).

**Figure 5:**
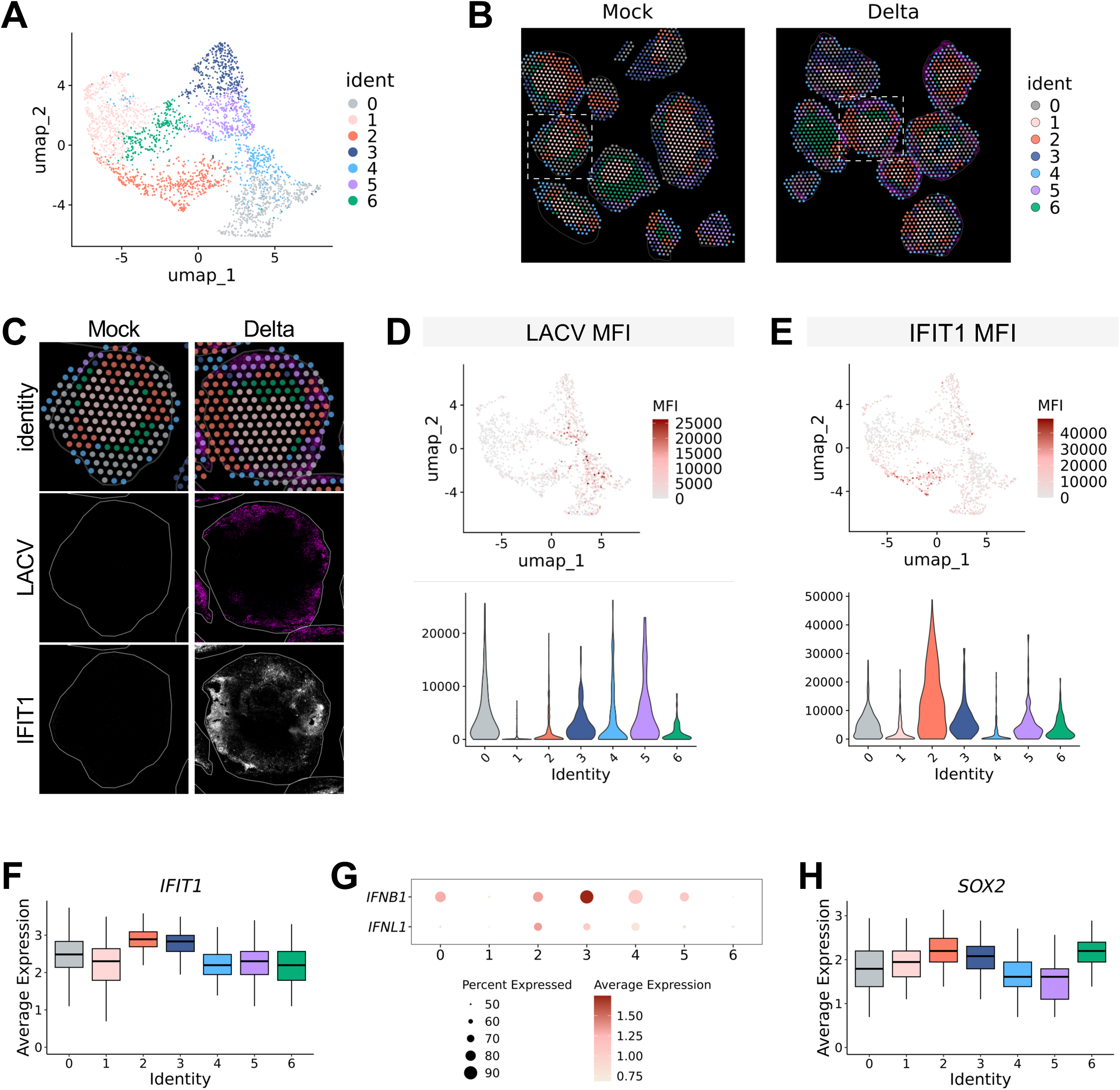
Spatial transcriptomics uncovers distinct regions of progenitor bystander activation. (A) Uniform manifold approximation and projection (UMAP) plot of spatial transcriptomics spots representative of 1407 spots in 8 ΔNSs-LACV infected organoids and 1470 spots in 9 mock-treated organoids. (B) Spatial clusters aligned to LACV-stained organoids in respective mock and ΔNSs-LACV-infected forebrain organoids. (C) Representative organoid (dashed white box in B) with aligned immunofluorescent images below; LACV glycoprotein (magenta) and IFIT1 (white). (D) Upper, UMAP plot and lower, violin plot of mean fluorescence intensity (MFI) of LACV signal within each spot in ΔNSs-LACV-infected organoids. (E) Upper, UMAP plot and lower, violin plot of MFI of IFIT1 signal within each spot in ΔNSs-LACV-infected organoids. (F) Box plot showing expression of *IFIT1* in ΔNSs-LACV-infected organoids. Bar represents mean and whiskers represent mean±SEM. (G) Dot plot showing variance-scaled expression of IFN beta (*IFNB1*) and IFN lambda (*IFNL1*) in ΔNSs-LACV-infected organoids. (H) Box plot showing expression of *SOX2* in ΔNSs-LACV-infected organoids. Bar represents mean and whiskers represent mean±SEM.

We aimed to then evaluate differential levels of infection across the seven clusters. To achieve this quantitatively, we first extracted spatially barcoded positions and integrated them with the corresponding immunofluorescence images (**Figure 5C, Supplemental Figure 4A and 4B**). Quantification of the mean fluorescence intensity (MFI) of LACV signal within each spot showed highly infected regions were represented by clusters 0, 3, 4 and 5 (**Figure 5D**); areas that primarily mapped to the edges of the organoids (**Figure 5C**). However, clusters 1, 2 and 6 had little to no infection (**Figure 5D**). Clusters 1 and 6 mapped primarily to the center of the organoids, where we do not typically see viral spread, unless type I IFN signaling is compromised (**Figure 4A**). Intriguingly, cluster 2 mapped along organoid edges, yet had little to no infection compared to other regions along the perimeter (**Figure 5B**). This suggests that cluster 2 might represent regions of intercellular communication capable of protective immunity.

Next, to investigate if either of these regions represented bystander activation, we similarly quantified MFI for IFIT1 protein expression within each spot (**Figure 5C**). Given the absence of IFIT1 signal in mock organoids, we focused further analysis on ΔNSs-LACV-infected organoids. We found that cluster 2 distinctly had the highest IFIT1 protein expression (**Figure 5E**). Although gene expression data showed all clusters were positive for IFIT1 transcript, clusters 2 and 3 had the highest expression (**Figure 5F**). Given we found that bystander activation was type I IFN-dependent (**Figure 4**), we evaluated gene expression of all type I IFNs and only identified IFNβ (*IFNB1*) as virally induced (**Figure 5G and Supplement Figure 4D**). Interestingly, we also found similar expression levels of IFN lambda (*IFNL1*), a type III IFN capable of inducing antiviral immunity (**Figure 5G**). Despite similar gene expression between clusters 2 and 3, these data suggest cluster 2 represents regions of bystander activation.

We previously noted that bystander activation was mediated by neuronal progenitors, thus, we sought to evaluate *SOX2* gene expression across clusters. We observed elevated *SOX2* expression in cluster 2; however other clusters also exhibited similar expression levels (**Figure 5H**). This indicates that not all intercellular communication of progenitors ultimately results in protective bystander antiviral responses. Taken together, by combining protein and gene expression dynamics, we uncovered distinct regions of progenitor bystander intercellular communication that elicit neuron intrinsic protective antiviral immunity.

### Spatial resolution reveals critical underpinnings of protective antiviral response in neurons

The identification of unique progenitor antiviral regions led us to further investigate the underlying mechanisms of protection following ΔNSs-LACV infection. Knowing cluster 2 had elevated IFIT1 expression and limited viral infection, we sought to identify clusters that lacked protection. Using the spot quantification of IFIT1 and LACV MFI (**Figure 5**), we examined correlations between IFIT1 and LACV protein expression. Similar to our earlier observation, cluster 2 demonstrates high IFIT1 protein expression and low LACV presence, further supporting its identity as a distinct region that can mount protective antiviral responses (**Figure 6A**). This stands in contrast to clusters 0 and 4, which had lower IFIT1 protein expression and elevated LACV signal.

**Figure 6:**
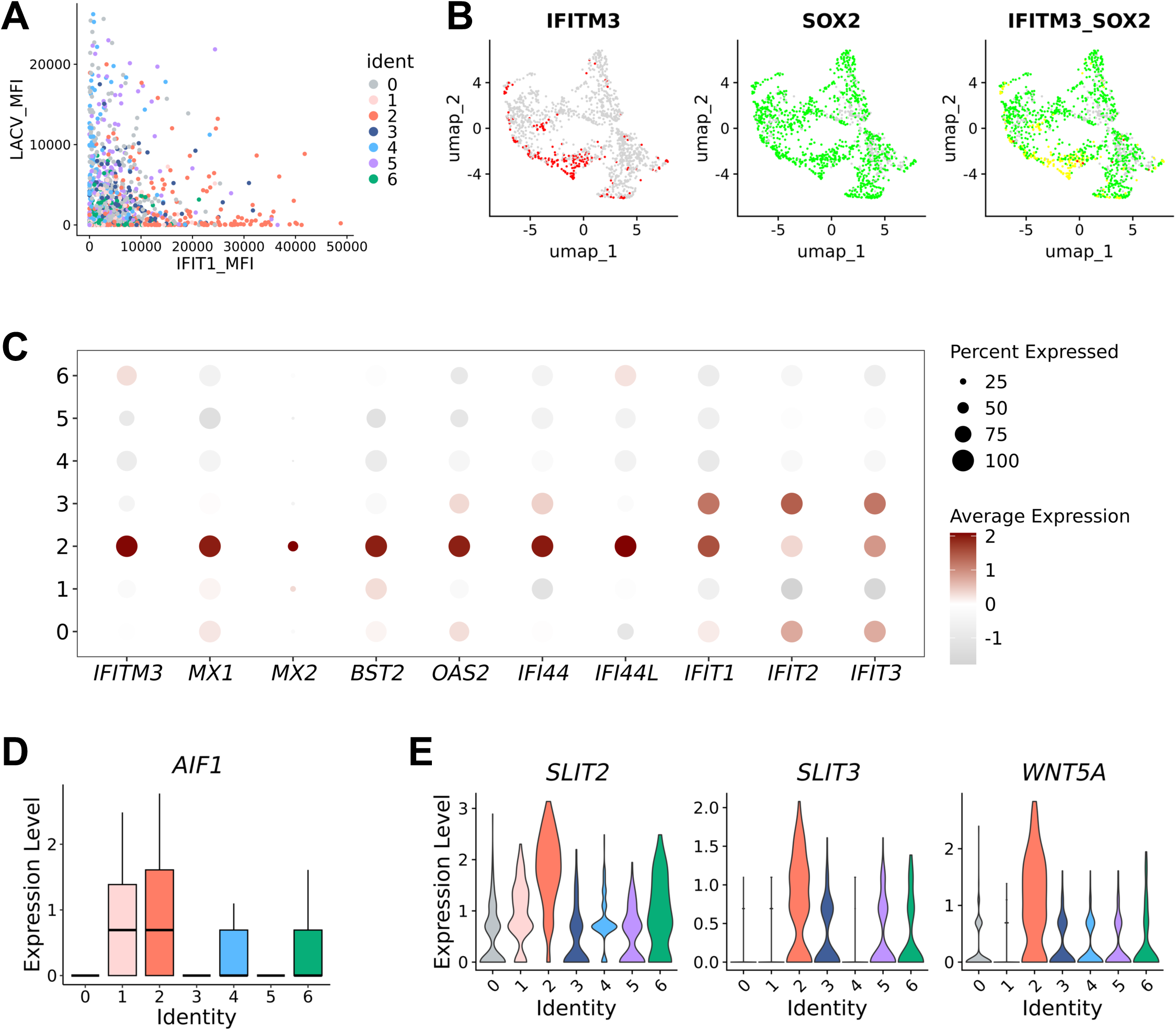
Spatial resolution reveals critical underpinnings of protective antiviral response in neurons. (A) Correlation of LACV and IFIT1 mean fluorescence intensity (MFI) in ΔNSs-LACV-infected organoids quantified in Figure 5. (B) Feature plots representing gene expression of *IFITM3* (red) or *SOX2* (green) in at least 50% of the populations. IFITM3_SOX2 represents colocalization of positive regions of ΔNSs-LACV-infected organoids. (C) Dot plot showing variance-scaled expression of interferon stimulated genes in ΔNSs-LACV-infected organoids. (D) Box plot showing expression of *AIF1* in ΔNSs-LACV-infected organoids. Bar represents mean and whiskers represent mean SEM. (E) Violin plot showing expression of *SLIT2*, *SLIT3*, and *WNT5A* in ΔNSs-LACV-infected organoids.

Next, we aimed to define the gene signatures contributing to this protective antiviral response. We conducted differential gene expression analysis comparing cluster 2 with clusters 0 and 4. This revealed *IFITM3*, a known inhibitor of LACV, as highly induced in cluster 2 (**Figure 6B**).^15^ Upon comparing this expression with *SOX2*, we observed colocalization between *IFITM3* and *SOX2*, further supporting a distinct neural progenitor antiviral region. Additionally, we identified other ISGs with established roles in LACV-antagonism, such as *MX1* and *MX2*, which were uniquely expressed in cluster 2 (**Figure 6C**). Intriguingly, we identified several ISGs in cluster 2 that had not been previously associated with LACV, including *BST2*, *OAS2*, and *IFI44*. Notably, IFI44L is a negative regulator of ISGs that was strongly upregulated in cluster 2. In contrast, similar to our observation with *IFIT1* expression, we observed broad expression of *IFIT2* and *IFIT3*. We also identified induction of *AIF1* which is a calcium-binding protein linked to the activation of microglia, a resident-brain macrophage (**Figure 6D**).^16^ This is particularly interesting as it suggests the potential for bystander neurons to communicate with other neighboring cells in the brain. These data suggest cluster 2 can uniquely orchestrate antiviral responses through robust induction of specific ISGs and putative communication with microglia.

We next sought to understand the underlying cellular properties that might confer cluster 2 with this unique antiviral capacity. We identified distinct expression of genes related to axonal guidance, specifically, repulsion factors *SLIT2*, *SLIT3* and *WNT5A* (**Figure 6E**).^17, 18^ Specifically, SLITs can guide neuronal pathfinding and migration.^19^ This suggests regions that are actively regulating axonal connection could be more protected from viral infection. Taken together, these results suggest potential underlying mechanisms of protection within unique progenitor antiviral regions where bystander progenitor neurons may protect themselves from infection while alarming neighboring cells.

## DISCUSSION

The innate immune interactions between uninfected bystander and infected neurons during viral infection remains incomplete. We designed this study to investigate the role of intercellular crosstalk in mediating intrinsic neuronal immunity and its contribution to protective antiviral responses. We found that in the absence of viral antagonism, neurons transcriptionally induce robust IFN signaling and can effectively signal to neighboring uninfected bystander neurons (**Figure 1**). Yet, in two-dimensional cultures, this dynamic response did not restrict viral spread. Interestingly, this differed in the context of viral infection in three-dimensional forebrain organoids, where we observed protective capacity (**Figure 3**). This finding underscores the importance of neuronal architecture and heterogeneity in facilitating intercellular communication to produce effective antiviral responses.

Previous work evaluating LACV infection in cerebral organoids found that committed neurons transcriptionally expressed fewer ISGs than neural progenitors^2^. Similarly, we observed the most robust induction of ISGs in neural progenitors, both transcriptionally and at the protein level. By considering location and infection status, we found that ISG-expressing neural progenitors were uninfected, bystander cells neighboring infected regions, indicating a previously unrecognized role. Of note, we uncovered the capacity for bystander neural progenitors to prevent viral replication and spread. However, application of spatial transcriptomics revealed that not all neuronal progenitors are the same, and specifically, that regionality was important (**Figure 5**). Removal of viral antagonism enabled us to unmask the intrinsic capacity of neuronal innate immune activation and signaling. We demonstrate that type I interferon was a contributor to protective neuronal intrinsic responses with pharmacological inhibition and genetic knockout (**Figure 4**). Yet, our results do not rule out a role for other inflammatory responses. In fact, we observed greater viral replication control with ruxolitinib treatment, a janus-kinase inhibitor that broadly limits innate immune signaling, as compared to knockout of the type I interferon receptor alone. Interestingly, our spatial transcriptomics data uncovered the induction of a type III interferon, *IFNL1*, suggesting a potential role in protection (**Figure 5**). Future studies are needed to directly address the role of type III interferons in neuronal intrinsic antiviral responses. Our work has added to the field by defining gene signatures of protective neuronal intrinsic responses using differential gene expression analysis between protected and infected regions. Of note, we found that protected regions expressed distinct antiviral genes, including ISGs previously found to protect against LACV infection.^15^ Future work should focus on determining the direct antiviral capacity of these ISGs in mediating protection against bunyaviral infection. Interestingly, investigation of protected regions uncovered enhanced expression of axonal repulsion factors^20^ making it intriguing to consider how induction of axonal repulsion factors might interface with intrinsic antiviral immunity. To test this model directly, future studies could focus on investigation with viruses that use different methods of viral spread.

Overall, our work reveals that intrinsic neuronal antiviral responses are intact within the context of cortical layers. Specifically, our work uncovers a previously unrecognized role for bystander neural progenitors in mediating signaling needed to orchestrate protective immunity amongst neurons. Further, we identify critical underpinnings of protection, including distinct antiviral genes. We envision that through defining underlying neuronal innate immune activation and signaling, we can leverage this foundational knowledge to inform development of novel antivirals to protect against neurotropic infection.

### Limitations of the study

One limitation to our study is the resolution of spatial transcriptomics where we cannot define gene expression changes at the individual cell level in relation to regional differences. Second, while our system allowed us to uncover neuronal intrinsic antiviral responses, it lacked other cell types of the brain, such as microglia, limiting our ability to evaluate neuronal communication with other cells.

## STAR METHODS

### KEY RESOURCES TABLE

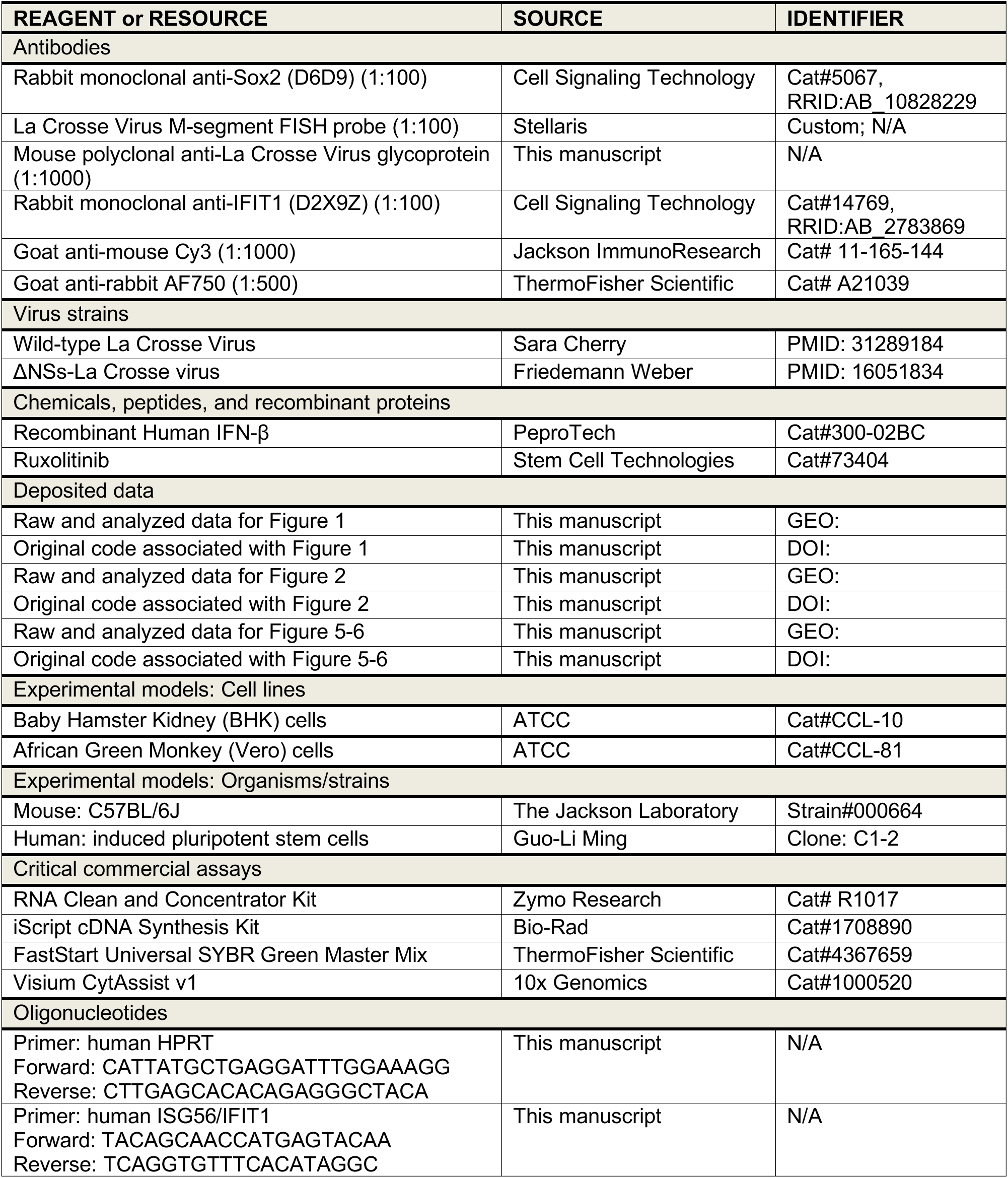
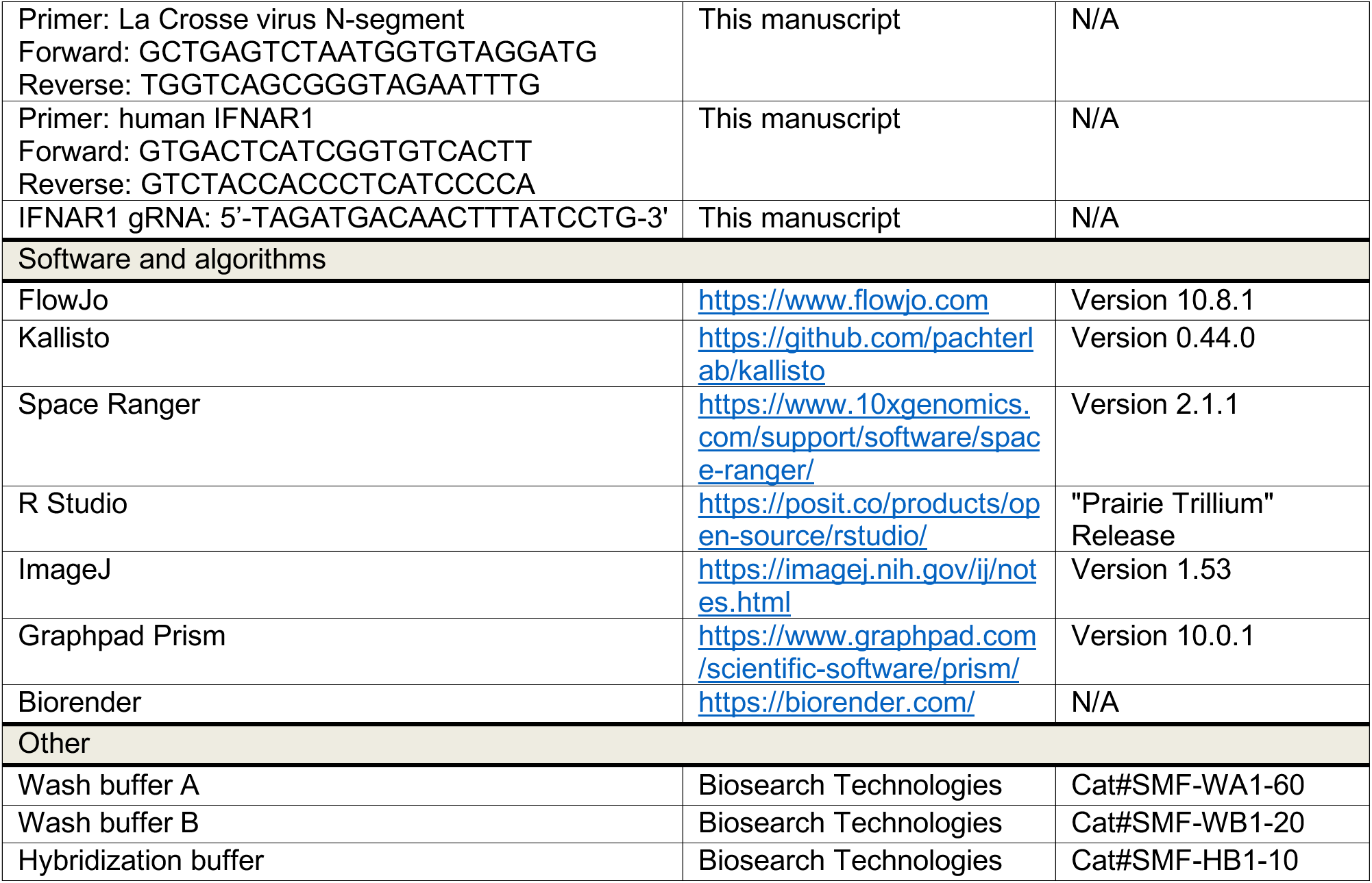

### RESOURCE AVAILABILITY

#### Lead contact

Further information and requests for resources and reagents should be directed to and will be fulfilled by the lead contact, Kellie A. Jurado.

#### Materials availability

Materials are available upon request from the lead author.

#### Data and code availability

Microscopy and sequencing data reported in this paper will be shared by the lead contact upon request. Any information required to reanalyze the data reported in this paper is available from the lead contact upon request.

### EXPERIMENTAL MODELS

Mice were used for cortical neuron isolation and bulk RNA-sequencing. Human induced-pluripotent stem cells were used to generate forebrain organoids. Further details are provided in the method details section.

#### Mice

C57BL/6J mice (The Jackson Laboratory, Strain#000664) were maintained at the University of Pennsylvania under specific pathogen free conditions, on a 12-hour light/dark cycle at 21C ± 1 C, 50% humidity ± 10%. All experiments were performed in adherence to the University of Pennsylvania’s approved IACUC protocol.

#### Cell lines

Baby hamster kidney (BHK) (ATCC, Cat#CCL-10) and African green monkey kidney (Vero) (ATCC, Cat#CCL-81) cells were grown in Dulbecco’s modification of eagle’s medium (Corning, Cat#10-013-CV) supplemented with 10% fetal bovine serum (Sigma-Aldrich, Cat#F2442), 1% penicillin-streptomycin (ThermoFisher Scientific, Cat#15140122), and 1% GlutaMAX (ThermoFisher Scientific, Cat#35050061). Human-induced pluripotent stem cell line C1-2 was kindly provided by Guo-Li Ming and maintained in mTSeR media (StemCell Technologies, Cat#100-1130) prior to forebrain organoid generation protocol. All cell lines were routinely tested and found to be Mycoplasma-free at the Cell Center Services Facility (University of Pennsylvania).

### METHOD DETAILS

#### Cortical neuron cultures

Embryos from embryonic day 16.5 of C57BL/6J mice (The Jackson Laboratory, Strain#000664) were dissected in cold 1x PBS. Cortices were resuspended and mechanically dissociated in room temperature Opti-MEM (Gibco, Cat#31985070) supplemented with 1% GlutaMAX. Cells were seeded on tissue culture treated plates coated with Poly-D-Lysine (Gibco, Cat#A3890401) according to manufacturer’s recommendations. OptiMEM-GlutaMAX was replaced with neuronal media consisting of Neurobasal (Invitrogen, Cat#21103049) supplemented with 1% penicillin/streptomycin, 1% GlutaMAX and 2% B-27 (Gibco, Cat#17504044). After 4-days *in-vitro* (DIV), neuronal media was supplemented with 0.5uM cytosine beta-D-arabinofuranoside (Sigma, Cat#C1768). Neurons were cultured at 37°C and 5% CO_2_ until 9-DIV where neuronal dendrites increased in number and branching.

#### Forebrain organoids

Forebrain organoids were generated using a previously described protocol.^21, 22^ Briefly, iPSCs were incubated at 5% CO_2_, 37C and maintained until 60-85% confluent. iPSCs exhibiting signs of differentiations were excluded from organoid generation. Additionally, only iPSCs between passage numbers 15 and 45 were used in the study. At 0-DIV, iPSCs were seeded in low-attachment 96-well plates (Corning, Cat#3474) at a density of 50,000 to 200,000 cells/well in mTeSR1 supplemented with 10µM of ROCK inhibitor (StemCell Technologies, Cat#72304). At 2-DIV, embryoid bodies (EB) were transferred using cut 1000uL pipette tips into 6-well plates in DMEM:F12 media (Gibco, Cat#11320033), supplemented with 20% Knockout Serum Replacement (Gibco, Cat#10828028), 1X GlutaMAX, 1X MEM Non-essential Amino Acids (Gibco, Cat#11140-050), 1X EmbryoMax 2-Mercaptoethanol (Sigma-Aldrich, Cat#ES-007-E), 1X Penicillin/Streptomycin, 1uM LDN193189 (StemCell Technologies, Cat#72147), 1μM SB-431542 (StemCell Technologies, Cat#72234) and 2ug/mL 0.2% heparin solution (StemCell Technologies, Cat#7980). Media changes were performed daily. At 6-DIV, EB media was replaced with induction media consisting of DMEM:F12, 1X N-2 Supplement (ThermoFisher Scientific, Cat#17502048), 1X Penicillin/Streptomycin, 1X NEAA, 1X GlutaMAX, 1 μM CHIR99021 (StemCell Technologies, Cat#72054), and 1 μM SB-431542. Round EBs with bright edges were selected at 7-DIV, coated with Matrigel, and plated on ultra-low-attachment 6-well plates (Corning, Cat#3471). Media was changed every other day until 14-DIV when EB-Matrigel complexes were disassociated. EBs, now organoids, were maintained until 35-DIV with daily media changes consisting of 1:1 Neurobasal media and DMEM/F12 supplemented with 1X N-2, 1X B-27, 1X GlutaMAX, 1X MEM Non-essential Amino Acids, 1X EmbryoMax 2-Mercaptoethanol, 1X Penicillin/Streptomycin and 2.5ug/mL of insulin (Sigma-Aldrich, Cat#I9278). All EBs and organoids were maintained at 5% CO_2_, 37C and incubated on a shaking platform at 120 revolutions per minute, except when EBs were complexed with Matrigel.

#### iPSC IFNAR1 knockout using CRISPR/Cas9

Human *IFNAR1* guide RNAs (gRNA) targeting early exons were selected from previous literature.^23^ gRNA and *S. pyogenes* Cas9 v2 were purchased from idtDNA (Cat#10007806). iPSCs displaying stem cell morphologies were selected to undergo electroporation. 300ng of gRNA was incubated at room temperature with 7.5μg of Cas9 for 10 minutes. Electroporation was then performed using Neon Electroporation Kit according to manufacturer’s guidelines (ThermoFisher Scientific, Cat#MPK1025). Briefly, 1×10^6^ iPSCs were resuspended in 5μL of Resuspension Buffer R. 5μL of resuspend cells was mixed and incubated with Cas9-gRNA complex. Using Neon pipette and tip, Cas9-gRNA was pulsed a single time into iPSCs at 1,200V for 30 milliseconds. Cells were immediately transferred into 24-well plate (Corning, Cat#3524) coated with Matrigel and containing mTeSR1 supplemented with 10μM ROCK inhibitor. Daily media changes were performed with mTeSR1 only until cells were 60% confluent and genomic DNA was extracted. IFNAR1 primers (Key Resource Table) targeting site of predicted gRNA-mediated Cas9 cut were designed and used to amplify region of interest. Amplicons were Sanger sequenced and clones with frameshift causing insertion/deletion mutations were maintained in culture. Clones underwent another round of subcloning to ensure single clonal population identity. These clones were again Sanger sequenced to confirm presence of frameshift causing deletion resulting in a premature stop codon in exon 2. The iPSC IFNAR1 knockout clone with no signs of differentiation was selected for validation and future experiments.

#### Virus stocks

Recombinant ΔNSs La Crosse Virus (ΔNSs-LACV) with a point-mutation in the non-structural protein of the S segment.^11^(Kindly provided by Friedemann Weber, Jutus-Liebig University, Germany) and wild-type La Crosse Virus (WT-LACV) (kindly provided by Sara Cherry) were propagated using the BHK cell line. For virus validation, RNA from infected BHK cells was reverse transcribed and a fragment of the LACV S segment was amplified by PCR as previously described.^11^ EcoRI digestion of RT-PCR products was used to distinguish WT-LACV (two 572 bp and 217 bp fragments) from the recombinant ΔNSs-LACV (silent mutation lacking EcoRI digestion site). TaiI digestion of the RT-PCR product was used to validate the point-mutation in recombinant ΔNSs-LACV (two fragments of 103 bp and 232 bp) from WT-LACV (no digestion site).

#### Virus infections

Murine cortical neurons were infected at 9-DIV by removing half of the media from cultures (neuron-conditioned media) and infecting the remaining volume at a multiplicity of infection (MOI) of 0.5 of WT- or ΔNSs-LACV at 37C. At 1-hour post-infection (HPI), infectious media was removed, and cells were replenished with neuron-conditioned media for collection at 16-HPI, unless otherwise noted. Vero cells were infected with 5 MOI ΔNSs-LACV and collected using 0.05% trypsin (Gibco, Cat#25300054) at 16-HPI for fluorescence in situ hybridization (FISH) probe validation. Human-derived forebrain organoids were placed in 12-well tissue culture treated plates with two organoids per well at 35-DIV. Culture media was completely removed and replaced with fresh culture media containing 2.5×10^6^ plaque forming units (PFU) of WT- or ΔNSs-LACV. Organoids were infected for 24 hours at 37°C and complete media changes occurred daily until 4-days post-infection (DPI).

#### Drugs and treatments

Human-derived forebrain organoids were infected with 2.5×10^6^ plaque forming units (PFU) of WT- or ΔNSs-LACV supplemented with 20uM ruxolitinib, a janus-kinase inhibitor (StemCell Technologies, Cat#73404) or 100 units/mL recombinant IFNβ (PeproTech, Cat#300-02BC) where indicated at 37°C for 24 hours. After infection, virus containing media was removed and replaced with fresh media supplemented with 20uM ruxolitinib or 100 units/mL recombinant IFNβ where indicated. Supplemented media changes were sustained until 4-DPI. To validate *IFNAR1* knockout in iPSCs and organoids, functional assays were performed on WT and IFNAR1 knockout iPSCs and 35-DIV forebrain organoids to determine response to IFNβ stimulation. Specifically, confluent iPSC cultures and forebrain organoids were treated with IFNβ for 6 hours prior to RNA extraction. Response to IFNβ was assessed by measuring ISGs *IFIT1, OAS2,* and *MX1* using RT-qPCR.

#### Neuron collection for flow cytometry

ΔNSs-LACV-infected and uninfected murine cortical neurons were collected at 16-HPI using an adapted previously described enzymatic and mechanical dissociation protocol.^24^ Briefly, 9-DIV murine cortical neurons were washed with 1x PBS warmed to 37C. Neurons were then treated with a warmed dissociation buffer containing accutase (StemPro, Cat#A1110501), papain (Worthington Biochemical Corporation, Cat#LK003176), EDTA (Invitrogen, Cat#15575-038) and 1x PBS at 37°C for 3-6 minutes until neuronal lifting was observed. Equal parts of neurobasal supplemented with 10% fetal bovine serum (Sigma-Aldrich, Cat#F2442) and 2mM EDTA were added to neurons to neutralize the dissociation. Neurons were then transferred to a conical, mechanically dissociated using a 10mL pipet and spun at 300xg at 4°C for 4 minutes. Supernatant was removed and fresh 3.7% PFA (ThermoFisher Scientific, Cat#J19943-K2) was added for 10 minutes at room temperature. Equal parts of FISH buffer containing 1xPBS and 0.2mg/mL RNase-free BSA (Sigma-Aldrich, Cat# B2518) was added and centrifuged at 300xg for 5 minutes at room temperature. Supernatant was removed and cells were resuspended in 500μL 70% ethanol then transferred to a 1.5mL Eppendorf.

#### Fluorescence in situ hybridization (FISH) staining and flow cytometry

Custom Stellaris® FISH Probes were designed against LACV M segment by utilizing the Stellaris® FISH Probe Designer (Biosearch Technologies, Inc.) available via Biosearch Technologies Stellaris designer resulting in 48 probes (**Supplemental Table 1**). Additionally, Stellaris® FISH Probes recognizing GAPDH or GFP (Biosearch Technologies, Inc., Cat#VSMF-3013-5 and Cat#VSMF-1017-5) were used. Fixed murine cortical neurons and Vero cells were hybridized with the LACV M segment, GAPDH, or GFP FISH Probe sets labeled with CAL Fluor Red 610, following the manufacturer’s instructions. Briefly, fixed cells were pelleted at 300xg for 5 minutes at room temperature and washed with wash buffer A (Biosearch Technologies, Inc., Cat# SMF-WA1-60). Cells were resuspended in hybridization buffer (Biosearch Technologies, Inc., Cat#SMF-HB1-10) containing the FISH Probe sets, then incubated in the dark at 37°C for 30 minutes. Cells were then pelleted and resuspended in wash buffer A for 30 minutes at 37°C. Cells were pelleted and incubated in wash buffer B (Biosearch Technologies, Inc., Cat#SMF-WB1-20) for 5 minutes at room temperature. Cells were then transferred to FACs tubes and washed with FISH buffer. Prior to flow cytometry, 1x DAPI was added to FISH probe-stained samples. All flow cytometry was conducted on the BD Biosciences LSRFortessa™ Cell Analyzer. All FACs was conducted on the BD Biosciences Influx sorter.

#### RNA extraction, cDNA generation and RT-qPCR

Cultured neurons or forebrain organoids were homogenized (MP Biomedical, Cat#116004500) in TRIzol (Invitrogen, Cat#15596018). RNA was extracted with Clean and Concentrator kit (Zymo, Cat#R1017) as per the manufacturer’s instructions and RNA concentrations were measured with the ThermoFisher Scientific Nanodrop One spectrophotometer. cDNA was generated using 1μg for neurons or 200ng for forebrain organoids of RNA and iScript cDNA synthesis kit (Bio-Rad, Cat#1708890) as per the manufacturer’s instructions. qPCR was performed with Power SYBR Green Master Mix (ThermoFisher Scientific, Cat#4367659). Reactions were run via QuantStudio3 (50C: 2’; 95C: 10’; 40x 95C: 15s, 60C: 1’) with the addition of a final melt curve (95C: 15s; 60C: 1’; 95C: 1’). All samples were loaded in technical duplicates. Melt curves were confirmed for each sample, and no-template controls were run to ensure no contamination. Average Ct value was calculated per sample, which was normalized against housekeeping (*hprt* or *HPRT*) expression. Normalized expression was presented relative to the appropriate control.

#### Bulk RNA-sequencing

RNA was processed for Bulk RNA-sequencing at the Children’s Hospital of Philadelphia High Throughput Sequencing Core. All data was analyzed using an adapted form of the open-source DIY transcriptomics lecture materials.^25^

#### Plaque assay

Infected supernatants from murine cortical neuron cultures or homogenates from forebrain organoids were serially diluted in DMEM (Corning, Cat#10-013-CV). Seeded BHK cells in a 6-well plate were treated with 200μL of diluted supernatants and incubated for 1 hour, with rotations every 15 minutes. Following incubation, inoculum was removed and replaced with MEM (Sigma-Aldrich, Cat#11430030) supplemented with 5% FBS, 1% GlutaMAX, 1% Non-essential amino acids, and 0.65% agarose (Lonza, Cat#50111). At 3-DPI, cells were fixed with 2mL 10% NBF (ThermoFisher Scientific, Cat#22050105) and visualized using 0.1% crystal violet (ThermoFisher Scientific, Cat#C581-25). Plaques were manually counted to calculate virus titer.

#### Immunofluorescence

Primary murine neurons were washed with 1x PBS and fixed with 4% PFA for 15 minutes at room temperature. For sectioned human forebrain organoids, samples were incubated in 4% PFA for 30 minutes at room temperature, while rocking, and incubated in 30% sucrose overnight before being embedded in OCT (Sakura, Cat#4583) and cryosectioned (5-10μm) using a Leica Cryostat. Wells and slides were washed three times with PBS prior to membrane disruption with PBS supplemented with Triton X (Sigma-Aldrich, Cat#T8787). Sample was blocked with PBS supplemented with 1% bovine serum albumin (Sigma-Aldrich, Cat# B2518) for 1 hour prior to overnight incubation with primary antibodies (Key Resources Table) diluted in blocking buffer. Samples were washed three times for 10 minutes with PBS supplement with 0.2% Tween20 (Biorad, Cat#1706531). After washing, samples were incubated for 1 hour in secondary antibodies (Key Resources Table) and 1xDAPI (ThermoFisher Scientific, Cat# AC202710100) diluted in blocking buffer. Following incubation, murine neuron samples were washed with PBST and imaged. Sectioned forebrain organoids were covered with mounting media and a coverslip was applied and sealed.

#### Spatial Transcriptomics

Mock-treated or ΔNSs-LACV-infected were sectioned (10μm) onto Visium CytAssist version 1 slides. Samples were stained with Eosin for processing with 10x CytAssist. Serial sections were stained with LACV glycoprotein targeting antibody, IFIT1 and DAPI for immunofluorescent imaging. Samples from CytAssist slides were processed for sequencing. Spatial data was aligned to LACV-stained organoids using 10X spaceranger. We next used Seurat to filter out spots with greater than 5% mitochondrial genes, excluding a total of 10 spots. To evaluate organoid complexity, we applied a principal component cut-off of 15 and a clustering resolution of 0.4. All remaining analysis was conducted using Seurat.

#### Data analysis and availability

All sequencing data was analyzed using R Studio. Graphs were plotted and statistical analyses were performed using GraphPad Prism. Data are expressed as Mean ± standard deviation (SD) or error of the mean (SEM). Number of biological samples used per experiment (n), number of individual experiments (N), and statistical tests used for each experiment are included in figure legends. Statistical significance was determined by Unpaired Student’s t tests for group means, One-Way or Two-Way ANOVA followed by post-Hoc multiple comparisons test as indicated. p<0.05 was considered as significant. ns = non-significant, *p<0.05, **p<0.01, ***p<0.001, ****p<0.0001. Flow cytometry plots were generated using FlowJo. Immunofluorescence images were visualized using NIS-Elements and ImageJ.

## ACKNOWLEDGMENTS

We thank Dr. Friedemann Weber from the Jutus-Liebig University, Germany, for providing the recombinant ΔNSs La Crosse virus; Dr. Sara Cherry from the University of Pennsylvania, Philadelphia, PA, USA, for La Crosse virus glycoprotein targeting antibodies; Molecular Pathology & Imaging Core (MPIC) from the University of Pennsylvania, Philadelphia, PA, USA, and 10x Genomics for assistance with spatial transcriptomics; Flow Cytometry Core facility at the University of Pennsylvania, Philadelphia, PA, USA, for sorting; Dr. Scott Sherrill-Mix from the University of Michigan from the University of Pennsylvania, Philadelphia, PA, USA, for assistance with sequencing analysis. We like to thank the MPIC Visium pilot grant, Penn Presidential PhD, Penn Presidential Faculty, and Laurie Breyer for funding to support the work.

## AUTHOR CONTRIBUTIONS

S.G.N.: study conceptualization, study design, experimental work, data interpretation, RNA-seq data and bioinformatic analysis and manuscript writing. C.V.: study design, experimental work, manuscript review and editing. C.B.: study design, experimental work, manuscript review and editing. G.M.: study design, data interpretation, and manuscript review. K.A.J.: funding acquisition, study conceptualization, study design, manuscript writing, supervision.

## SUPPLEMENTAL INFORMATION

**Supplemental Figure 1:**
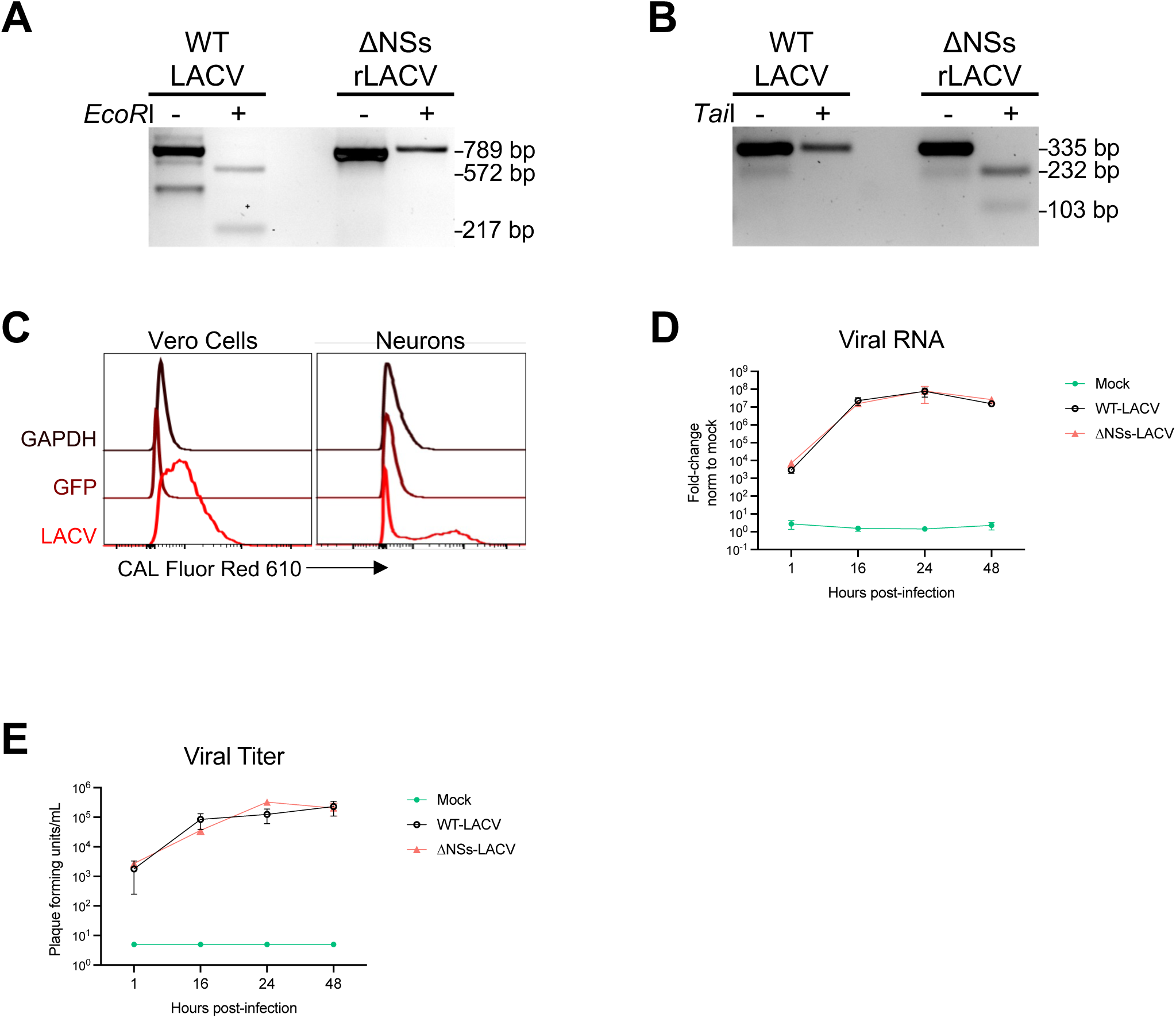
Recombinant virus and FISH probe validation. (A-B) BHK cells were infected with wild-type La Crosse Virus (WT-LACV) or recombinant ΔNSs La Crosse Virus (ΔNSs-LACV) at 5 MOI and RNA was extracted at 16-hours post-infection (HPI). Lack of EcoRI digestion of ΔNSs-LACV demonstrates the silent mutation used to distinguish recombinant virus (A). TaiI digestion validates ΔNSs-LACV’s inability to express NSs due to the ablation of the reading-frame as in (B). 1 experiment. (C-D) Fold change of LACV RNA relative to *Hprt* housekeeping gene determined by quantitative RT-PCR (C) and virion production determined by plaque assay (D) kinetics following WT- or ΔNSs-LACV infection of murine cortical neurons at 1-, 16-, 24-, and 48-HPI. Neurons pooled from 1 litter per independent experiment. N=3. Mean±SEM. (E) Vero cells and primary cortical neurons infected with 5 MOI and 0.5 MOI recombinant ΔNSs La Crosse Virus (ΔNSs-LACV), respectively. Cells were collected for FISH staining at 16 hours post-infection with probes targeting GAPDH (positive control), GFP (negative control) or LACV (M segment). Histograms represent cells analyzed using flow cytometry.

**Supplemental Figure 2:**
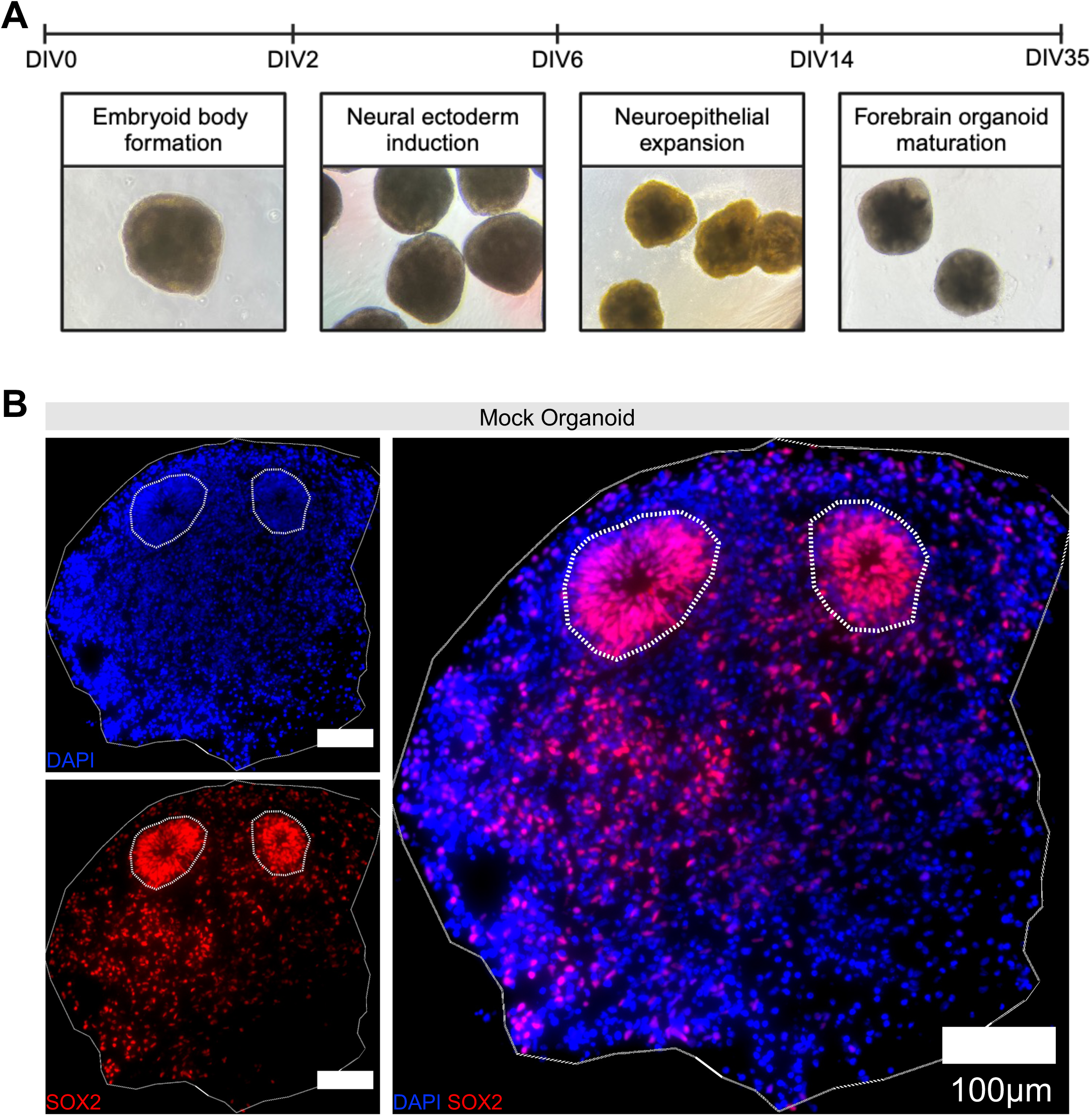
Radial organization and progenitor cells in forebrain organoids. (A) Model and brightfield images of 35-DIV forebrain organoid generation. (B) Immunofluorescent staining with progenitor marker SOX2 (red) and DAPI (blue) to demarcate neural rosettes. Scale bars: 100µm. Solid lines outline organoids and dotted lines outline neural rosettes based on DAPI and SOX2 staining.

**Supplemental Figure 3:**
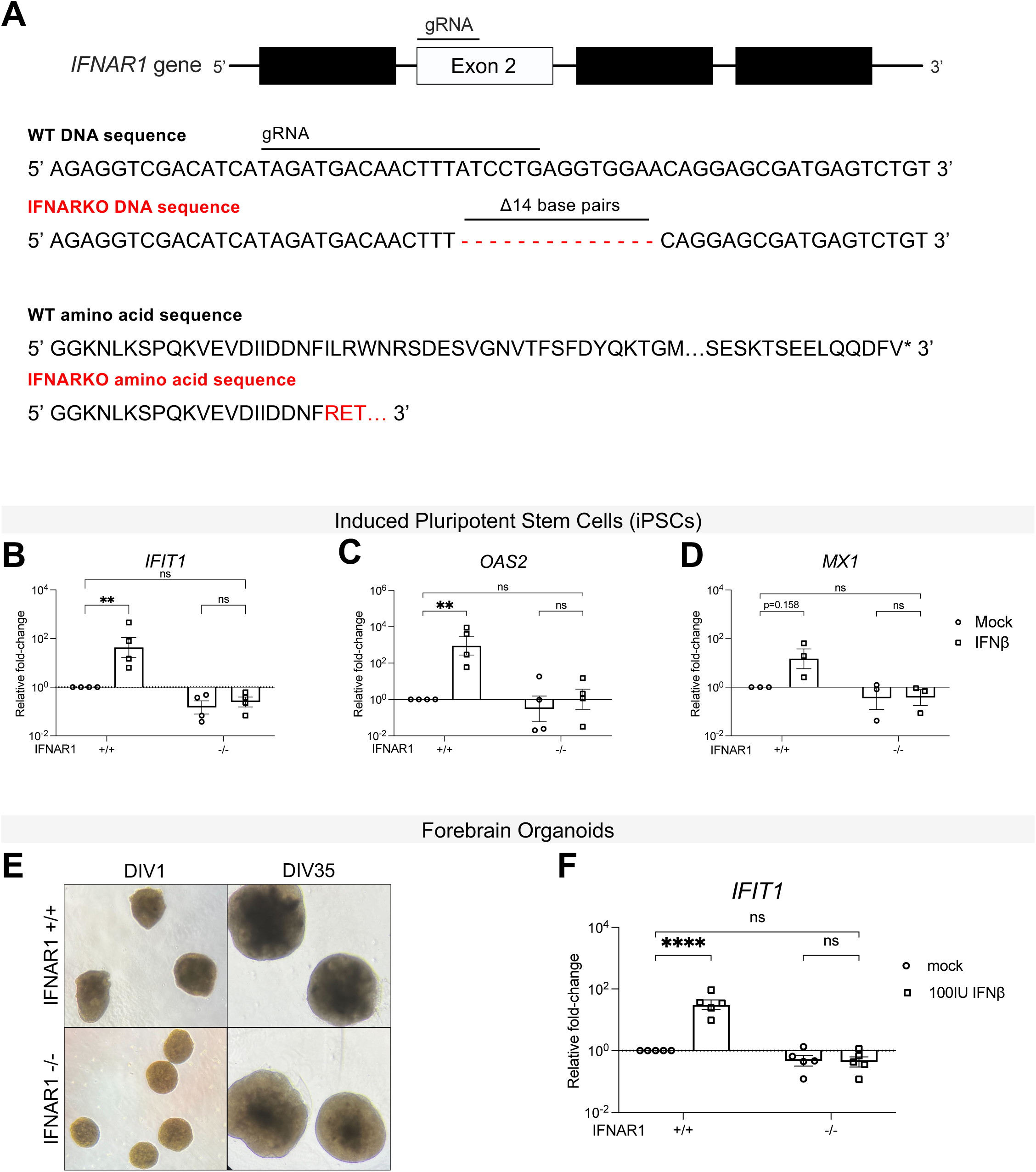
IFNAR knockout forebrain organoids do not respond to IFNβ stimulation. (A) Schematic of CRISPR/Cas9 guideRNA-mediated knockout of *IFNAR1* gene. (B-D) Wild type and IFNAR1 knockout iPSCs treated with 100 units/mL recombinant IFNβ or mock-treated for 6 hours. Fold change of *IFIT1* (B), *OAS2* (C), and *MX1* (D) RNA as determined by quantitative RT-PCR relative to the housekeeping gene *HPRT*, normalized to mock. N=4. Mean ± SEM. Two-way ANOVA followed by Tukey’s test. (E) Representative images of WT or IFNAR1 knockout embryoid bodies (DIV1) and forebrain organoids (DIV35). (F) WT and IFNAR1 knockout 35-DIV forebrain organoids treated with 100 units/mL recombinant IFNβ or mock-treated for 6 hours. Fold change of *IFIT1* RNA as determined by quantitative RT-PCR relative to the housekeeping gene *HPRT*, normalized to mock. N=5. Mean ± SEM. Two-way ANOVA followed by Tukey’s test, ns p>0.05, *p<0.05, **p<0.01 ****p<0.0001.

**Supplemental Figure 4:**
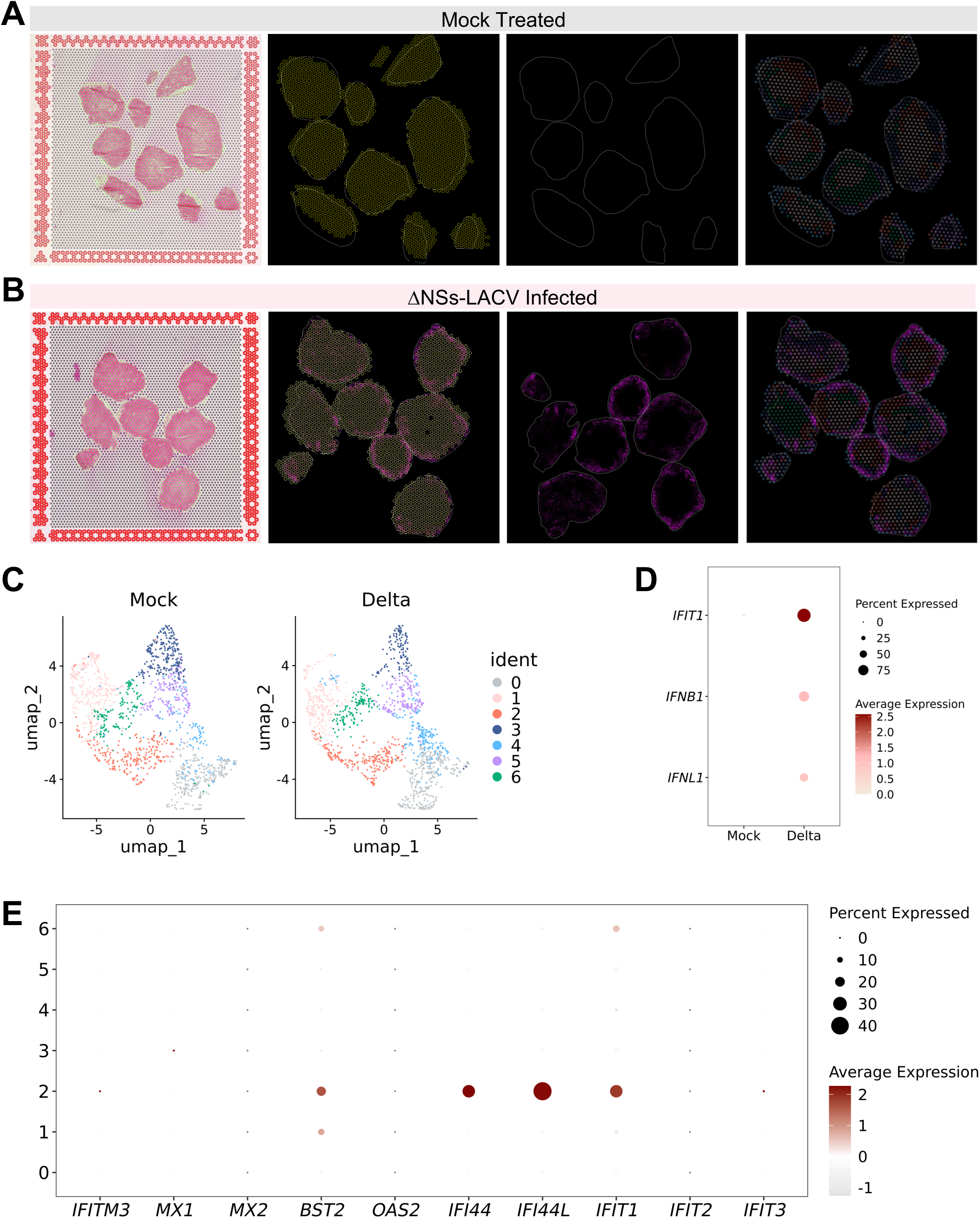
Spatial transcriptomics alignment and clustering reveal minimal basal ISG expression in forebrain organoids. (A-B) Mock-treated (A) or ΔNSs-LACV-infected (B) forebrain organoids with eosin staining aligned with fiducial markers; LACV (magenta) immunofluorescence staining aligned with 10x spatial spots; LACV (magenta) immunofluorescence images alone; and LACV (magenta) immunofluorescence images overlayed with UMAP clustering of spots. (C) UMAP clustering of spots grouped by conditions mock-treated (left) or ΔNSs-LACV-infected (right). (D) Dot plot of *IFIT1*, *IFNB1*, *IFNL1* normalized and variance-scaled gene expression, and percent spots expressing, grouped by conditions.

## REFERENCES

1. Cho, H. et al. Differential innate immune response programs in neuronal subtypes determine susceptibility to infection in the brain by positive-stranded RNA viruses. Nat. Med. 19, 458– 464 (2013).

2. Winkler, C. W. et al. Neuronal maturation reduces the type I IFN response to orthobunyavirus infection and leads to increased apoptosis of human neurons. J. Neuroinflammation 16, 229 (2019).

3. Griffin, D. E. Immune responses to RNA-virus infections of the CNS. Nat. Rev. Immunol. 3, 493–502 (2003).

4. Delhaye, S. et al. Neurons produce type I interferon during viral encephalitis. Proc. Natl. Acad. Sci. U. S. A. 103, 7835–7840 (2006).

5. Chakraborty, S., Nazmi, A., Dutta, K. & Basu, A. Neurons under viral attack: Victims or warriors? Neurochem. Int. 56, 727–735 (2010).

6. Lampron, A., ElAli, A. & Rivest, S. Innate Immunity in the CNS: Redefining the Relationship between the CNS and Its Environment. Neuron 78, 214–232 (2013).

7. Klein, R. S. et al. Neuroinflammation During RNA Viral Infections. Annu. Rev. Immunol. 37, 73–95 (2019).

8. Blakqori, G. et al. La Crosse bunyavirus nonstructural protein NSs serves to suppress the type I interferon system of mammalian hosts. J. Virol. 81, 4991–4999 (2007).

9. Verbruggen, P. et al. Interferon Antagonist NSs of La Crosse Virus Triggers a DNA Damage Response-like Degradation of Transcribing RNA Polymerase II. J. Biol. Chem. 286, 3681– 3692 (2011).

10. Sorgeloos, F., Kreit, M., Hermant, P., Lardinois, C. & Michiels, T. Antiviral Type I and Type III Interferon Responses in the Central Nervous System. Viruses 5, 834–857 (2013).

11. Blakqori, G. & Weber, F. Efficient cDNA-based rescue of La Crosse bunyaviruses expressing or lacking the nonstructural protein NSs. J. Virol. 79, 10420–10428 (2005).

12. Krishnaswami, S. R. et al. Using single nuclei for RNA-seq to capture the transcriptome of postmortem neurons. Nat. Protoc. 11, 499–524 (2016).

13. Amamoto, R. et al. Probe-Seq enables transcriptional profiling of specific cell types from heterogeneous tissue by RNA-based isolation. eLife 8, e51452 (2019).

14. Williams, C. G., Lee, H. J., Asatsuma, T., Vento-Tormo, R. & Haque, A. An introduction to spatial transcriptomics for biomedical research. Genome Med. 14, 68 (2022).

15. Basu, R. et al. Identification of age-specific gene regulators of La Crosse virus neuroinvasion and pathogenesis. Nat. Commun. 14, 2836 (2023).

16. Schwab, J. M. et al. AIF-1 expression defines a proliferating and alert microglial/macrophage phenotype following spinal cord injury in rats. J. Neuroimmunol. 119, 214–222 (2001).

17. Li, L., Hutchins, B. I. & Kalil, K. Wnt5a Induces Simultaneous Cortical Axon Outgrowth and Repulsive Axon Guidance through Distinct Signaling Mechanisms. J. Neurosci. 29, 5873– 5883 (2009).

18. Gonda, Y., Namba, T. & Hanashima, C. Beyond Axon Guidance: Roles of Slit-Robo Signaling in Neocortical Formation. Front. Cell Dev. Biol. 8, (2020).

19. Wong, K., Park, H. T., Wu, J. Y. & Rao, Y. Slit proteins: molecular guidance cues for cells ranging from neurons to leukocytes. Curr. Opin. Genet. Dev. 12, 583–591 (2002).

20. Gillet, J. P., Derer, P. & Tsiang, H. Axonal transport of rabies virus in the central nervous system of the rat. J. Neuropathol. Exp. Neurol. 45, 619–634 (1986).

21. Chiang, C.-H. et al. Integration-free induced pluripotent stem cells derived from schizophrenia patients with a DISC1 mutation. Mol. Psychiatry 16, 358–360 (2011).

22. Lancaster, M. A. et al. Cerebral organoids model human brain development and microcephaly. Nature 501, 373–379 (2013).

23. Wang, R., Yang, J. F., Senay, T. E., Liu, W. & You, J. Characterization of the Impact of Merkel Cell Polyomavirus-Induced Interferon Signaling on Viral Infection. J. Virol. 97, e01907–22.

24. Jerber, J., Haldane, J., Steer, J., Pearce, D. & Patel, M. Dissociation of neuronal culture to single cells for scRNA-seq (10x Genomics). (2020).

25. Berry, A. S. F. et al. An Open-Source Toolkit To Expand Bioinformatics Training in Infectious Diseases. mBio 12, 10.1128/mbio.01214-21 (2021).

