## Supplemental Table 1 for "Neuronal interactions in forebrain organoids lead to protective antiviral responses"

| Sequence Name | Sequence | Fluorescent Modification |
|---|---|---|
| LACV M Segment _1 | gcaatcatgccaatcagaga | CAL Fluor Red 610 |
| LACV M Segment _2 | aggtcatcaccaacttctat | CAL Fluor Red 610 |
| LACV M Segment _3 | gtacaatctgcgctgcaaac | CAL Fluor Red 610 |
| LACV M Segment _4 | ttgtggtatctgtctgaagc | CAL Fluor Red 610 |
| LACV M Segment _5 | aggtgttcgcaagtttgatc | CAL Fluor Red 610 |
| LACV M Segment _6 | caggcatggaactgtacaga | CAL Fluor Red 610 |
| LACV M Segment _7 | ccacactctgtgaatggatg | CAL Fluor Red 610 |
| LACV M Segment _8 | gctcttaggcttttataacc | CAL Fluor Red 610 |
| LACV M Segment _9 | taagaccagtaccgcagtaa | CAL Fluor Red 610 |
| LACV M Segment _10 | ctctcctaaaaccatggagt | CAL Fluor Red 610 |
| LACV M Segment _11 | catttccaacatgtcgtctg | CAL Fluor Red 610 |
| LACV M Segment _12 | cccgatcaacaatccaatga | CAL Fluor Red 610 |
| LACV M Segment _13 | cagcatcttcatattgacca | CAL Fluor Red 610 |
| LACV M Segment _14 | ttataggttttcctgtgagc | CAL Fluor Red 610 |
| LACV M Segment _15 | aagttcatcatccactttgc | CAL Fluor Red 610 |
| LACV M Segment _16 | gaggttcaaaaatggcccag | CAL Fluor Red 610 |
| LACV M Segment _17 | tcttttgttgcttttggcaa | CAL Fluor Red 610 |
| LACV M Segment _18 | ccctttaactgagttgcaat | CAL Fluor Red 610 |
| LACV M Segment _19 | atccctgttattatagggac | CAL Fluor Red 610 |
| LACV M Segment _20 | aaagcacccgcaatatcatc | CAL Fluor Red 610 |
| LACV M Segment _21 | aagtacactctccatgttgt | CAL Fluor Red 610 |
| LACV M Segment _22 | agctgttagggctagattta | CAL Fluor Red 610 |
| LACV M Segment _23 | ccttcaacaatgcattacct | CAL Fluor Red 610 |
| LACV M Segment _24 | gattggaatacaccacctga | CAL Fluor Red 610 |
| LACV M Segment _25 | ggcctcaaattcttctagac | CAL Fluor Red 610 |
| LACV M Segment _26 | ttcccaacatttggatttct | CAL Fluor Red 610 |
| LACV M Segment _27 | tcttggatgagctttacgtc | CAL Fluor Red 610 |
| LACV M Segment _28 | agttcccagaatctttcatt | CAL Fluor Red 610 |
| LACV M Segment _29 | ccatctgttagttgcaacat | CAL Fluor Red 610 |
| LACV M Segment _30 | ttaatagggtaccggacagt | CAL Fluor Red 610 |
| LACV M Segment _31 | cctcaatattgtgcacatct | CAL Fluor Red 610 |
| LACV M Segment _32 | gatgtgcggcaagttttttg | CAL Fluor Red 610 |
| LACV M Segment _33 | tatgcactctctataccttc | CAL Fluor Red 610 |
| LACV M Segment _34 | atgcaggtacttctgattca | CAL Fluor Red 610 |
| LACV M Segment _35 | ggccagttgagtacaatttt | CAL Fluor Red 610 |
| LACV M Segment _36 | acccaacctgatgattgata | CAL Fluor Red 610 |
| LACV M Segment _37 | gatccaaatacacacccatc | CAL Fluor Red 610 |
| LACV M Segment _38 | ctgtaggtctctacagtttt | CAL Fluor Red 610 |
| LACV M Segment _39 | ccattgccataaatagttcc | CAL Fluor Red 610 |
| LACV M Segment _40 | gcatgcttggtaatcattgt | CAL Fluor Red 610 |
| LACV M Segment _41 | gaagcagttaatacagccgg | CAL Fluor Red 610 |
| LACV M Segment _42 | caattgtggtgtgcaacgtt | CAL Fluor Red 610 |

| | | |
|---|---|---|
| LACV M Segment _43 | ggagtcacaagaatcctgtc | CAL Fluor Red 610 |
| LACV M Segment _44 | ctgtgcacaccattttcaat | CAL Fluor Red 610 |
| LACV M Segment _45 | ttaattgtgagtgtgttccc | CAL Fluor Red 610 |
| LACV M Segment _46 | gctagttctatgataggctt | CAL Fluor Red 610 |
| LACV M Segment _47 | tccaagttttacacctttca | CAL Fluor Red 610 |
| LACV M Segment _48 | atgcatcttcatgctttcta | CAL Fluor Red 610 |
